# Coordinated beak–tongue mechanics enable dexterous seed manipulation in songbirds

**DOI:** 10.64898/2026.04.22.720076

**Authors:** Maja Mielke, Falk Mielke, Nicholas W. Gladman, Dan A. Tatulescu, Anthony Herrel, Coen P. H. Elemans, Sam Van Wassenbergh

## Abstract

Dexterous manipulation of objects relies on precise coordination between anatomical elements. In seed-eating birds, seeds are manipulated and dehusked using both the beak and tongue, but the functional roles and coordination of these structures remain unresolved. Here, we quantified the 3D movements of the upper beak, lower beak, tongue, and seed in a hard-biting and a weak-biting songbird species using X-ray Reconstruction of Moving Morphology (XROMM) and measured contractile properties of their primary jaw muscles. We show that the tongue serves as the main tool for seed rotation, transport, and stabilization. Multi-dimensional, high-frequency movements of the upper and lower beaks reveal that efficient seed processing depends on high mobility of the kinetic avian skull. Strong and weak biters differ in feeding kinematics and jaw muscle speeds, suggesting ecological specialization of cranial mechanics. The complexity, precision, and tight coordination of beak and tongue motions show that the avian cranium rivals the dexterity of the primate hand despite limited degrees of freedom.

## Introduction

For millions of years, birds and their ancestors have relied on seeds as their principal or complementary food source. Seeds are highly nutritious, energy-rich, and durable, and they have played an essential role in the success of the avian lineage in the course of bird evolution (Larson et al., 2016). The ability to feed on seeds helped avian dinosaurs survive the end-Cretaceous extinction event 66 million years ago (Larson et al., 2016), and since then, birds have evolved into more than 11,000 species. Today, more than 600 bird species are granivorous, i.e., feeding predominantly on seeds (Tobias et al., 2022). Thus, the ability to eat seeds was and remains critical to the evolution and survival of extant birds.

While large birds swallow seeds whole, smaller birds face a vital, yet extremely challenging, task when feeding on seeds. They need to split and remove the indigestible husk before swallowing a seed’s inner kernel (Kear, 1962; Meij & Bout, 2006; Mielke & Van Wassenbergh, 2022; Nuijens & Zweers, 1997; Van der Meij et al., 2004; Ziswiler, 1965). While some mammals use their dexterous hands for such complex behaviors that require skill and precision (Iwaniuk & Whishaw, 1999; Padberg et al., 2007; Pouydebat et al., 2014; Sustaita et al., 2013), songbirds seemingly manage this feat using their beaks alone. Precise seed positioning and stabilization are essential for successful dehusking (Meij & Bout, 2006; Ziswiler, 1965): birds must position the seed carefully between the fairly sharp ridges of the upper and lower beaks while preventing it from falling out during force application. Previous observations have proposed a role of the tongue for seed positioning and medial stabilization (Ziswiler, 1965; Zweers et al., 1994), but so far, we lack both conclusive evidence for involvement of the tongue and quantitative data on tongue and seed movements during seed manipulation.

Efficient seed processing involves three-dimensional mobility of the jaws that goes far beyond a mere open-close movement (e.g., Mielke & Van Wassenbergh, 2022). The kinetic skull of birds in principle allows for such complex 3D movements of the upper and lower beaks. The lower beak has multiple degrees of freedom due to its multi-joint suspension from the skull via the quadrate bones, which allows for medio-lateral movements during seed processing (Dawson et al., 2011; Meij & Bout, 2006; Mielke & Van Wassenbergh, 2022; Nuijens & Zweers, 1997; Ziswiler, 1965). Furthermore, most birds utilize some degree of cranial kinesis, i.e., movement of the upper beak relative to the head, during feeding (Bock, 1964, 1966; Bout & Zweers, 2001; Estrella & Masero, 2007; Heuvel, 1992; Mielke & Van Wassenbergh, 2022; Zusi, 1984). However, we lack quantification of the complex 3D movements of the upper and lower beaks and how they are coordinated with seed and tongue movements during seed manipulation.

Evolutionary diversification of beak morphology and bite force has profound implications for songbird feeding performance (Meij & Bout, 2006; Van der Meij et al., 2004; Ziswiler, 1965). Yet, their functional consequences for feeding mechanics remain poorly understood. As shown prominently in Darwin’s finches, beak morphology is adapted to feeding on specific food items, which may differ in size, shape, and hardness (Grant & Grant, 2006; Herrel et al., 2010; Schluter & Grant, 1984). Beak size and bite force are under strong selective pressure and can change within a few generations in bird populations in response to changes in the abundance of certain food types (Diamant & Yeh, 2025; Grant & Grant, 1995), a phenomenon that promotes rapid songbird evolution (Podos & Schroeder, 2024). Selection for high bite force may give a granivorous bird access to seeds of particular hardness but may impede the maximal velocity of beak movement, confining the abilities of a hard-biting songbird to produce fast bird song (Herrel et al., 2009). However, we don’t know if, and how, hard and weak biters also differ in their jaw muscle performance, feeding kinematics, and husking performance.

The primary goal of this study was to determine how granivorous songbirds solve the challenge of seed husking using intricate feeding mechanics. We quantified the three-dimensional movements of the beak, tongue, and seed during seed processing in feeding songbirds using X-Ray Reconstruction of Moving Morphology (XROMM, Figure 1). We establish that tongue motion is crucial to position, rotate, and stabilize seeds. Furthermore, we show that seed processing requires fast, precisely coordinated, and complex movements of both the tongue and the entire kinetic skull. Our insights reveal how the avian cranium operates as an integrated functional system with precision and agility that virtually serves as a bird’s surrogate “hand”.

**Figure 1:**
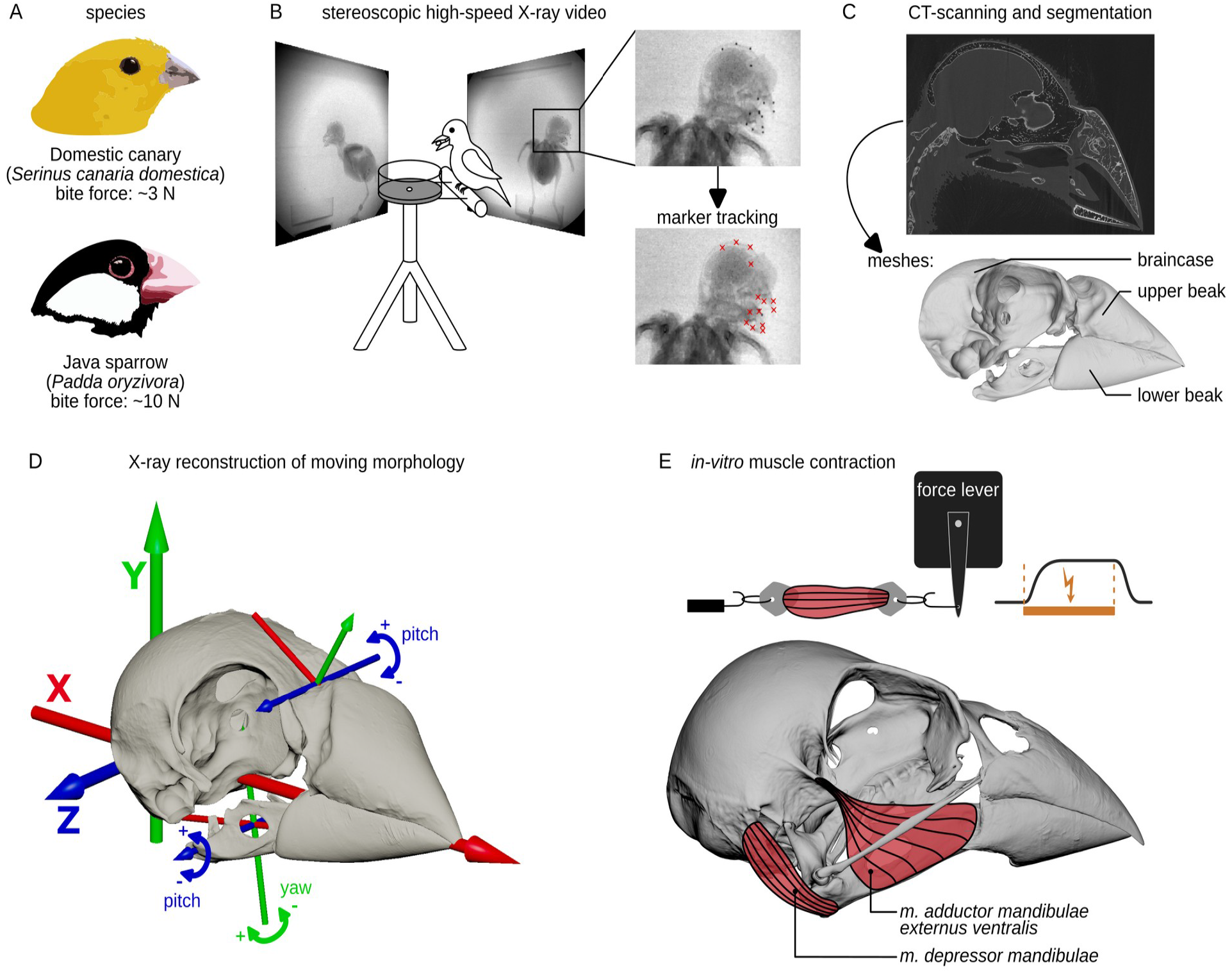
Summary of methods used in this study. **A)** Study species: the weak-biting domestic canary and the hard-biting Java sparrow. **B)** Schematic visualization of the biplanar high-speed X-ray videography. Marker positions were tracked in XMALab. **C)** Micro-CT-scans of the bird skulls and segmentation of rigid bodies, here represented on the skull of a Java sparrow. **D)** X-ray reconstruction of Moving Morphology (XROMM). Visualization of the anatomical coordinate system (thick axes) and the joint coordinate systems of upper and lower beaks (thin axes), here represented on the skull of a Java sparrow. For kinematic analyses, we quantified pitch angles of upper and lower beaks and yaw angles of the lower beak relative to the braincase. **E)** Measurements of contraction and relaxation rates of beak opener muscle (*m. depressor mandibulae*) and beak closer muscle (*m. adductor mandibulae externus ventralis*), here represented on the skull of a canary.

Furthermore, we tested whether specialization for high bite force is associated with distinct cranial kinematics and jaw muscle speed by comparing husking behavior and muscle performance in a hard-biting and a weak-biting songbird species, the domestic canary (*Serinus canaria*) and the Java sparrow (*Padda oryzivora,* Figure 1). We show that these species differ in beak-tongue kinematics and jaw muscle contractile properties, which suggests that ecological speciation drives distinct cranial mechanics, motor skills, and feeding techniques.

## Results

### The role of the tongue in seed transport and stabilization

To determine if, and how, the tongue contributes to transport and stabilization of seeds, we quantified the three-dimensional movements of the tongue and seed via X-ray markers in songbirds feeding on hemp seeds (see Methods). Birds process seeds in three distinct phases between pickup and swallowing (Mielke & Van Wassenbergh, 2022): (i) positioning (seed translations and rotations), (ii) biting (cracking the husk), and (iii) dehusking (removing the husk). Each of these phases involves its own typical movement patterns of the upper beak, lower beak, and tongue (Appendix 1—figure 1 and Appendix 1—figure 2). For analyzing the role of the tongue in seed manipulation, we focused on the positioning and biting phases, as seed integrity is not guaranteed during the dehusking phase, and therefore the seed’s X-ray marker may no longer reliably represent seed position. We quantified the correlation of seed and tongue translations by measuring the skewness of the distribution of correlation values in a permutation test (see Materials and Methods for details). Positive and negative skew values indicate negative and positive correlation, respectively.

**Figure 2:**
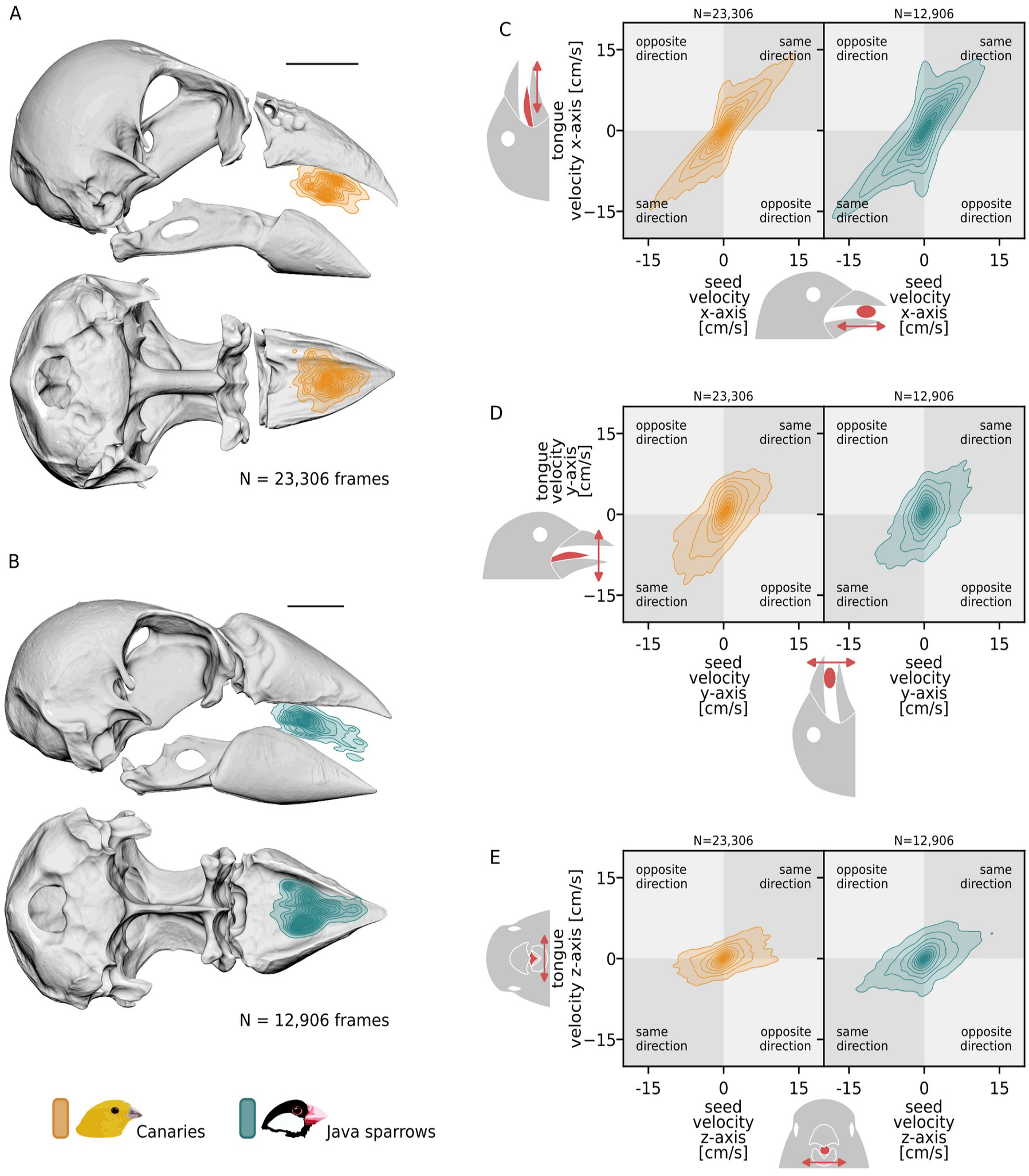
The tongue is critical for seed transport during positioning. Distributions of seed locations during positioning show that the seed is moved considerably within the beak (**A** and **B**). Relationships of tongue and seed velocities (**C**-**E**) indicate that the tongue and seed move mostly together in the same direction in both species, especially along the antero-posterior axis (C). All plots show kernel density estimates with darker colors indicating more frequent occurrences. Only data of ’positioning’ phases are shown. Skull images shown in lateral and ventral views, with the lower jaw hidden in the latter. Scale bars in A and B: 5 mm. In C-E, upper right and lower left quadrants indicate movements of seed and tongue marker in the same direction along the given axis (positive correlation). Upper left and lower right quadrants indicate opposing movements along the given axis (negative correlation). Red elements in icons in C-E indicate the object (tongue or seed) and axis of the anatomical coordinate system (x: posterior-to-anterior axis, y: ventral-to-dorsal axis, z: left-to-right axis). N denotes the number of frames included for calculation of kernel density estimates. All plots show the pooled data of two individuals per species.

Our data establish that during positioning, the birds used their tongue to move the seed within the beak to bring it into the correct position for a biting attempt. The location of the seed covered the majority of the antero-posterior (X-axis) and medio-lateral (Z-axis) range of the beak during that phase (Figure 2 A+B). Specifically, 95% of antero-posterior seed translation occurred within a range of 4.0 mm in canaries and 6.7 mm in Java sparrows. For the medio-lateral (Z-)axis, 95% of seed locations covered a range of 3.1 mm and 3.8 mm in canaries and Java sparrows, respectively. A high positive correlation of tongue velocity and seed velocity, especially along the antero-posterior axis (skew between -1.69 and -1.56 in canaries and between -1.55 and -1.41 in Java sparrows), shows that the tongue is crucial for transporting the seed within the beak during this phase (Figure 2 C-E, Appendix 1—table 1, Appendix 1—figure 3).

**Figure 3:**
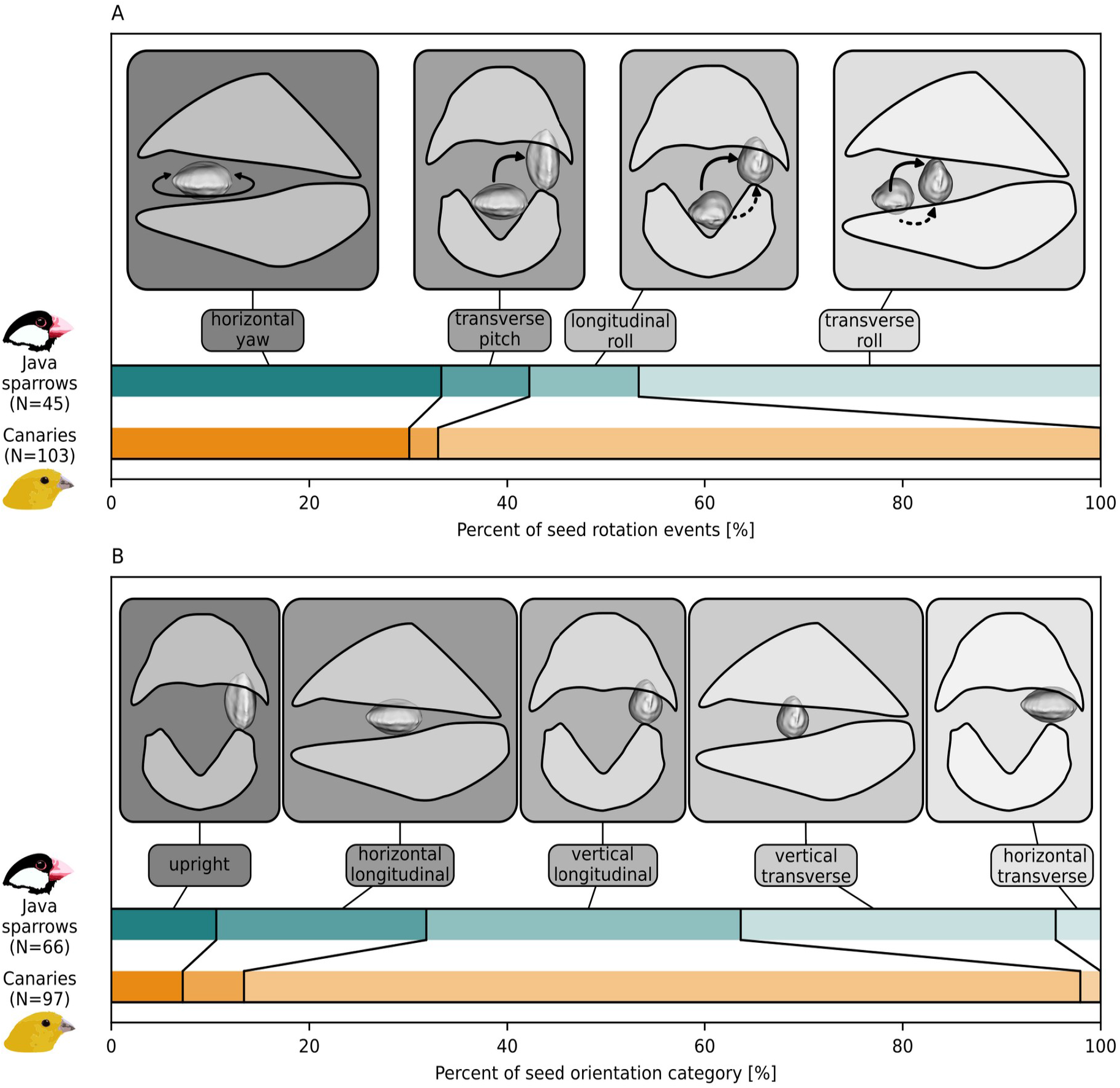
Birds use distinct types of seed rotation during positioning and seed orientation during biting. **A**) Four dominant types of seed rotation in canaries and Java sparrows. Solid arrows in (A) indicate approximate typical trajectories of seeds during rotation, dashed arrows indicate alternative options of trajectories that the birds used only occasionally. **B**) Five dominant types of seed orientation during biting in canaries and Java sparrows. ’Upright’ refers to a special case of transverse orientation, with the long axis of the seed lying parallel to the sagittal plane of the beak. In A+B, ‘longitudinal’ and ‘transverse’ refer to the long axis of the seed being parallel and orthogonal to the long axis of the beak, respectively. Seed rotations and orientations are visualized on only one side for clarity. Seeds and beaks are not drawn to scale. N denotes the sample size (number of analyzed seed rotation or biting events). Wide boxes show beaks in lateral view, narrow boxes show beaks in frontal view. All plots show the pooled data of two individuals per species.

During the positioning phase, the birds furthermore use their tongues to perform complex seed rotations to bring the seed into the correct orientation for a biting attempt. In canaries, 78% of analyzed positioning events included seed rotations; in Java sparrows, this was 48%. The birds used four different kinds of seed rotation: horizontal yaw, transverse pitch, longitudinal roll, and transverse roll (Figure 3A, Video 1). Seed rotation was characterized by fast antero-posterior movements of the tongue in canaries but less so in Java sparrows (Appendix 1—figure 4).

**Figure 4:**
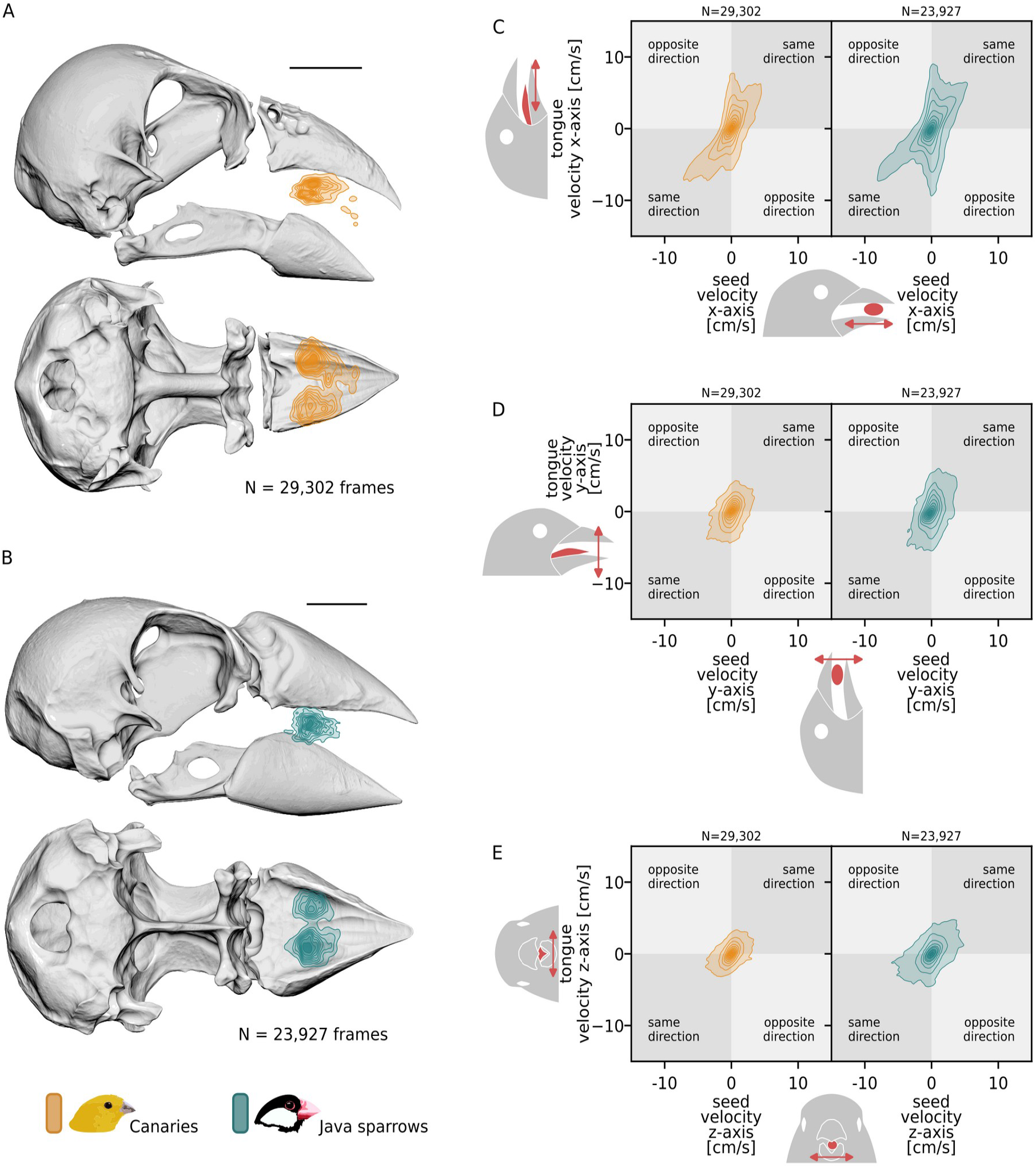
The tongue is critical for seed stabilization during biting. Distributions of seed locations during biting indicate the consistent seed position in either of the lateral grooves of the upper palate (**A** and **B**). In both species, the tongue and seed move less compared to the positioning phase (**C-E**, cf. Figure 2), indicating a stabilizing function of the tongue. All plots show kernel density estimates with darker colors indicating more frequent occurrences. Only data of ’biting’ phases are shown. Skull images shown in lateral and ventral views, with the lower jaw hidden in the latter. Scale bars in A and B: 5 mm. In C-E, upper right and lower left quadrants indicate movements of seed and tongue marker in the same direction along the given axis (positive correlation). Upper left and lower right quadrants indicate opposing movements along the given axis (negative correlation). Icons in C-E indicate the object (tongue or seed) and axis of the anatomical coordinate system (x: posterior-to-anterior axis, y: ventral-to-dorsal axis, z: left-to-right axis). N denotes the number of frames included for calculation of kernel density estimates. All plots show the pooled data of two individuals per species.

During biting, the seed is fixed between the upper and lower beaks, and the bird applies adductive forces to crack open the shell. In contrast to Java sparrows, canaries had a strong preference for a ‘vertical-longitudinal’ seed orientation during biting (82/97 biting events, Figure 3B). Independent of seed orientation, the birds positioned seeds in either of the two lateral grooves in their upper palate (Figure 4 A+B). Compared to positioning, the measured velocities of both seed and tongue were much lower, suggesting that the tongue stabilized the seed during biting (Figure 4 C-E). The correlation of seed and tongue movements was lower than during positioning, possibly due to the more static nature of the biting phase (Appendix 1—figure 5, Appendix 1—table 1).

**Figure 5:**
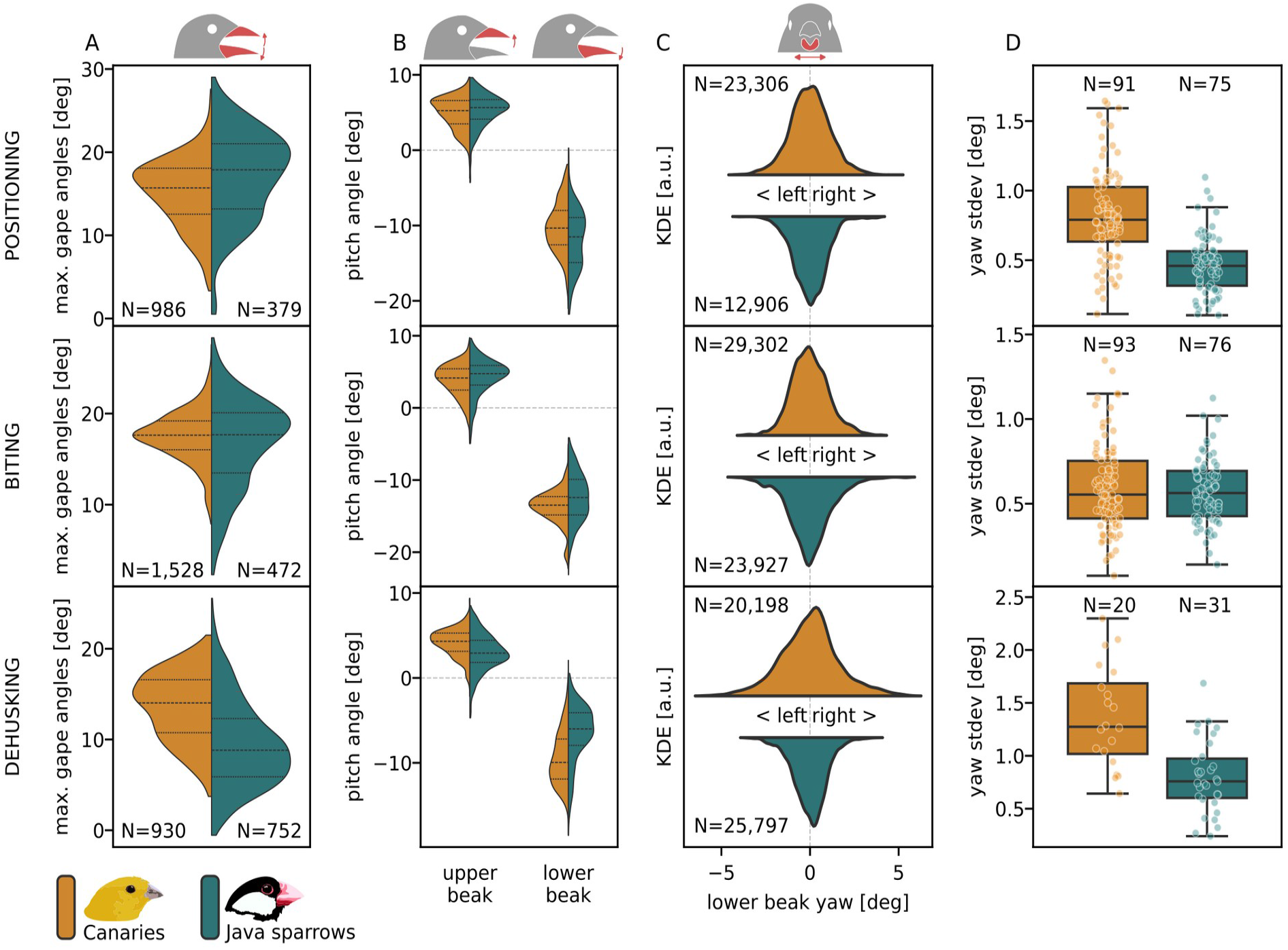
Motions of both the upper and the lower beaks are involved in seed processing. Distributions of peaks in total gape angle reveal differences between phase types and species **(A)**. Both upper and lower beaks contribute to total gapes **(B)**. Kernel density estimates (KDE) of yaw angles show that both species use medio-lateral rotations of the lower beak during all phases **(C).** Standard deviations of yaw angles of lower beak reveal differences between the species for the positioning and dehusking phase **(D)**. Dashed lines in A and B and boxplots in D indicate the quartiles of the distributions. The whiskers in D indicate the highest and lowest data point that lies within the 1.5 inter-quartile range of the upper and lower quartile. N denotes sample sizes (A+B: number of extracted peaks in total gape angle, C: number of frames included for calculation of kernel density estimates, D: number of sequences of which yaw stdev was calculated). All plots show the pooled data of two individuals per species. See Appendix 1—table 2 and Appendix 1—table 3 for results of the statistical analyses. **Source data 1.** Data of maximal total gape angles (A) and associated pitch angles of upper and lower beaks (B). **Source data 2.** Standard deviations of lower beak yaw rotations (D).

In conclusion, the tongue plays a crucial role in the precise positioning, manipulation, and cracking of seeds.

### Cranial kinesis and maximal beak opening

In small songbirds, processing large seeds requires considerably wide beak gapes. Upper beak rotations (cranial kinesis) may help achieve such wide gapes while keeping lower jaw depression at moderate levels. However, we don’t know how much the upper and lower beaks contribute to beak gape angle during seed processing. We analyzed the beak’s pitch angles to quantify how the widest necessary beak openings vary among the different phases of seed processing and how much the upper and lower beaks contribute to total beak gape. First, we detected the peaks of the total gape angle over time (Figure 5A). Then, we extracted the pitch angles of the upper and lower beaks for each frame with such a peak in total gape angle (Figure 5B).

In general, the maximal total gape reached up to 28 degrees in canaries and 29 degrees in Java sparrows (Figure 5A). The lower beak contributed up to 22 degrees in canaries and 23 degrees in Java sparrows (Figure 5B). The upper beak contributed up to 10 degrees to total gape in both species (Figure 5B), indicating extensive use of cranial kinesis.

The total gape during local maxima was relatively similar between the species during positioning and biting events (Figure 5A). For these phase types, the differences in total gape between the species averages were significant but small (below 1.7 degrees, Appendix 1—table 2). This is in line with the mean pitch angles of the upper and lower beaks not differing more than 1.4 degrees between the species during positioning and biting.

During dehusking, canaries used wider total gapes than Java sparrows (canaries: 13.6±0.1 degrees, Java sparrows: 9.9±0.1 degrees, difference between the species: 3.6±0.2 degrees, 94%hdi_can-jav_=[3.3, 4.0]). This is based on a wider opening in both the upper and lower beaks in canaries compared to Java sparrows (4.1±0.1 vs. 3.1±0.1 degrees for the upper beak, -9.4±0.1 vs. -6.8±0.1 degrees for the lower beak, Figure 5B, Appendix 1—table 2).

Taken together, we conclude that upper beak rotations (cranial kinesis) contribute considerably to beak opening during all seed processing phases. Furthermore, the maximal pitch angles during seed dehusking can differ between species, which suggests that birds use species-specific strategies in beak usage during husk removal.

### Multi-dimensional movements of the lower beak

In contrast to the upper beak, the lower beak is able to move in the medio-lateral direction, i.e., sideways (Meij & Bout, 2006; Mielke & Van Wassenbergh, 2022). To investigate the involvement of such movements in seed processing, we quantified the range of medio-lateral rotation in feeding songbirds. We measured the yaw angles of the joint coordinate system (JCS) of the lower beak (the JCS representing a virtual joint between the lower beak and braincase). To quantify the extent of yaw, we measured the standard deviation of the distributions of used yaw angles that we gained from the XROMM data for the different phases of seed processing. The higher the standard deviation, the wider the range of lower beak yaw angles.

Both species used medio-lateral movements of the lower beak but to different extents, depending on the phase type (Figure 5 C+D, Video 2). During biting, the range of yaw rotation was similar between the two species (Δstdev=0.03±0.04 degrees, 94%hdi_can-jav_=[-0.04, 0.10], Figure 5D, Appendix 1—table 3). Compared to Java sparrows, canaries showed a significantly wider range of yaw rotations during positioning (Δstdev=0.35±0.04 degrees, 94%hdi_can-jav_=[0.27, 0.44]) and during dehusking (Δstdev=0.67±0.12 degrees, 94%hdi_can-jav_=[0.43, 0.89], Figure 5D). Both species showed the widest range of mandible yaw during dehusking, which suggests that this phase of the feeding cycle demands particularly high beak agility.

The differences in mandible yaw rotation during dehusking reflect differences in the technique of husk removal between the species. To separate the husk from the kernel, canaries inserted the edge of the lower beak in the crack of the husk and widened the gap with rotational movements of the mandible (Video 3). By doing so, canaries achieved splitting the husk in two halves along the edge of the seed and removing these two halves while leaving the inner kernel intact (20/20 dehusking events). In contrast, Java sparrows frequently crushed the inner kernel and/or husk by continuing to bite on the seed while attempting to separate the husk from the kernel (15/28 dehusking events, Video 3). We conclude that the ability for wide yaw rotations may be beneficial for controlled husk removal.

The suspension of the lower jaw via the quadrate bones allows not only for medio-lateral rotations but also for longitudinal fore/aft translations, which may aid in splitting the husk (‘slicing’) during the biting phase. However, the birds tested here scarcely used slicing; such translations occurred within a range of only 1.2 mm in canaries and 0.6 mm in Java sparrows (measured as the range between the 3% and 97% percentiles of antero-posterior translations of the JCS of the lower beak during biting phases). This was not more than the translation during positioning and dehusking, indicating that these movements likely had no functional use specifically for opening the husk.

We conclude that medio-lateral rotations rather than longitudinal translations of the lower beak play a key role in seed manipulation. Such rotations may differ in their extent among phase types and species and may benefit successful husking of seeds.

### Coordination of upper and lower beak rotations

To maintain control while manipulating a seed, birds may benefit from some degree of independent (low-correlation) movements of the upper and lower beaks. To assess the degree of correlation in beak movements, we analyzed the combination of angular velocities of the upper and lower beak pitch angles. Angular velocity quantifies the direction and degree of change in pitch angle over time. Positive values indicate increasing angles (elevation); negative values indicate decreasing angles (depression). For each frame of each phase type, we quantified the angular velocities of the upper and lower beaks (Figure 6). If both values are positive, the upper and lower beaks move upwards together. If both are negative, they move downwards together. Both cases indicate a positive correlation. If one value is positive and the other negative, the upper and lower beaks move in opposite directions, i.e., with negative correlation.

**Figure 6:**
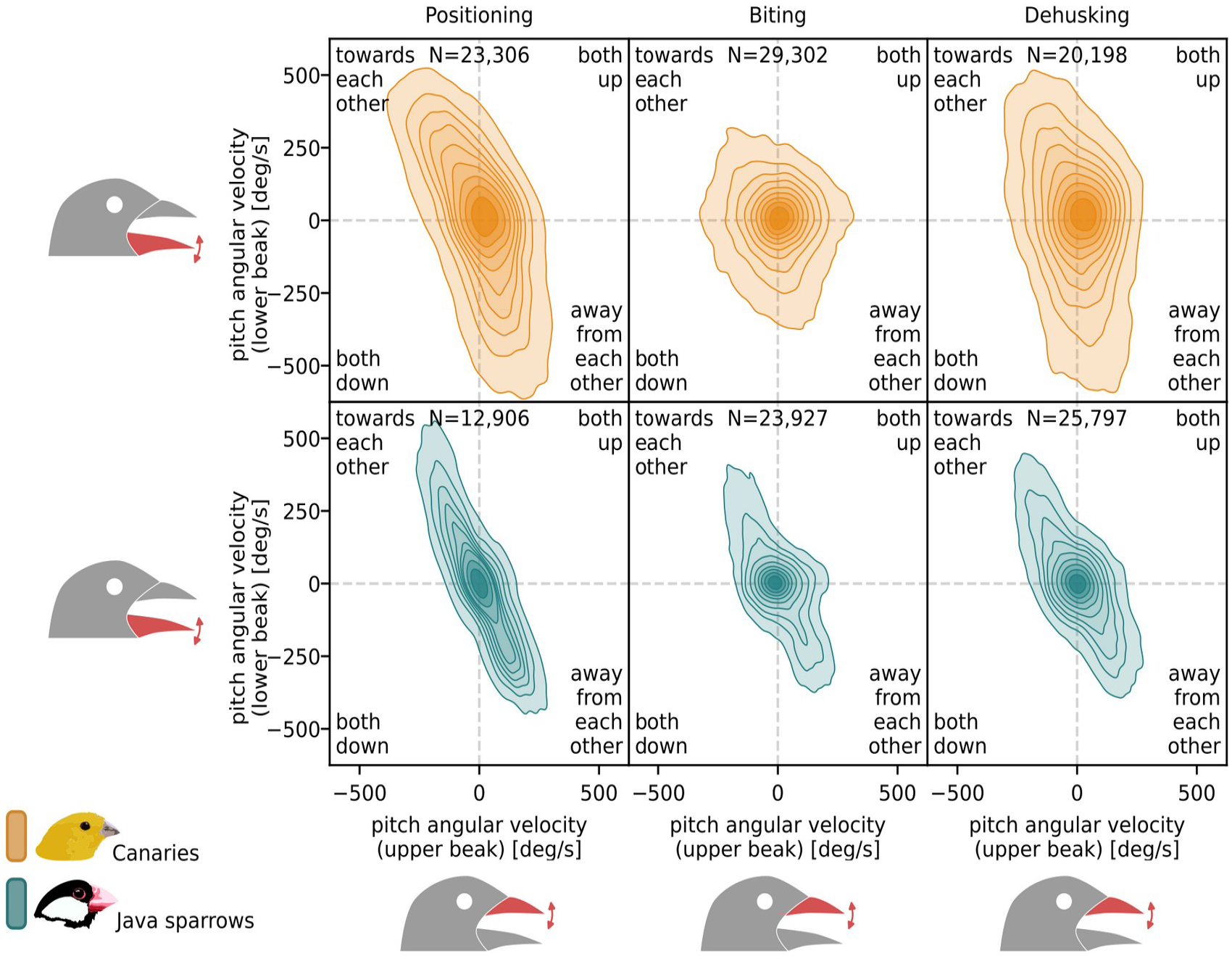
Coordination of upper and lower beak movements during positioning, biting, and dehusking. The narrow and diagonal density plot in the left column indicates strong negative correlation of upper and lower beaks during positioning. See main text for details. Plots show kernel density estimates of pitch angular velocities of the upper and lower beaks during positioning, biting, and dehusking. More frequent combinations of values are indicated by darker colors. N denotes the number of frames included for calculation of kernel density estimates. All plots show the pooled data of two individuals per species.

Our data revealed that the coordination of the upper and lower beaks pitch rotations differed with phase type and species (Figure 6, Appendix 1—table 4, Appendix 1—figure 6). We quantified the correlation of upper and lower beak correlation by measuring the skewness of the distribution of correlation values in a permutation test (see Materials and Methods for details). Positive and negative skew values indicate negative and positive correlation, respectively. Both species showed the most pronounced opposing movements of the upper and lower beaks during seed positioning (canaries: skew between 0.79 and 0.96; Java sparrows: skew between 1.19 and 1.74). During biting, both species, but especially canaries, used relatively more movement of the upper and lower beaks in the same direction. While skew values even indicate slight positive correlation in canaries (skew between -0.28 and -0.08), Java sparrows retain negative correlation, albeit at lower degrees compared to the positioning phase (skew between 0.61 and 0.69). During dehusking, we observed more opposing movements again, with skew values indicating negative correlation (canaries: skew between 0.23 and 0.36; Java sparrows: skew between 0.76 and 1.34). Independent of the phase type, the upper and lower beaks moved in opposite directions more strictly in Java sparrows than in canaries.

Taken together, our results indicate that (i) seed positioning and seed dehusking involve the highest degrees of negatively correlated movements of the upper and lower beaks, and (ii) canaries utilize more independent movements of the upper and lower beaks than Java sparrows.

### Coordination of beak movement and seed transport

Precise manipulation of the seed during the positioning phase requires good coordination of beak opening and seed transport. To explore how the birds coordinate movements of the beak with transport of the seed during that phase, we analyzed the relationship of seed translations with beak gape and beak angular velocity (Figure 7). During seed positioning, the seed is mainly transported along the antero-posterior axis via movements of the tongue (see Figure 2C). Hence, we took seed velocity along this axis into consideration in this analysis.

**Figure 7:**
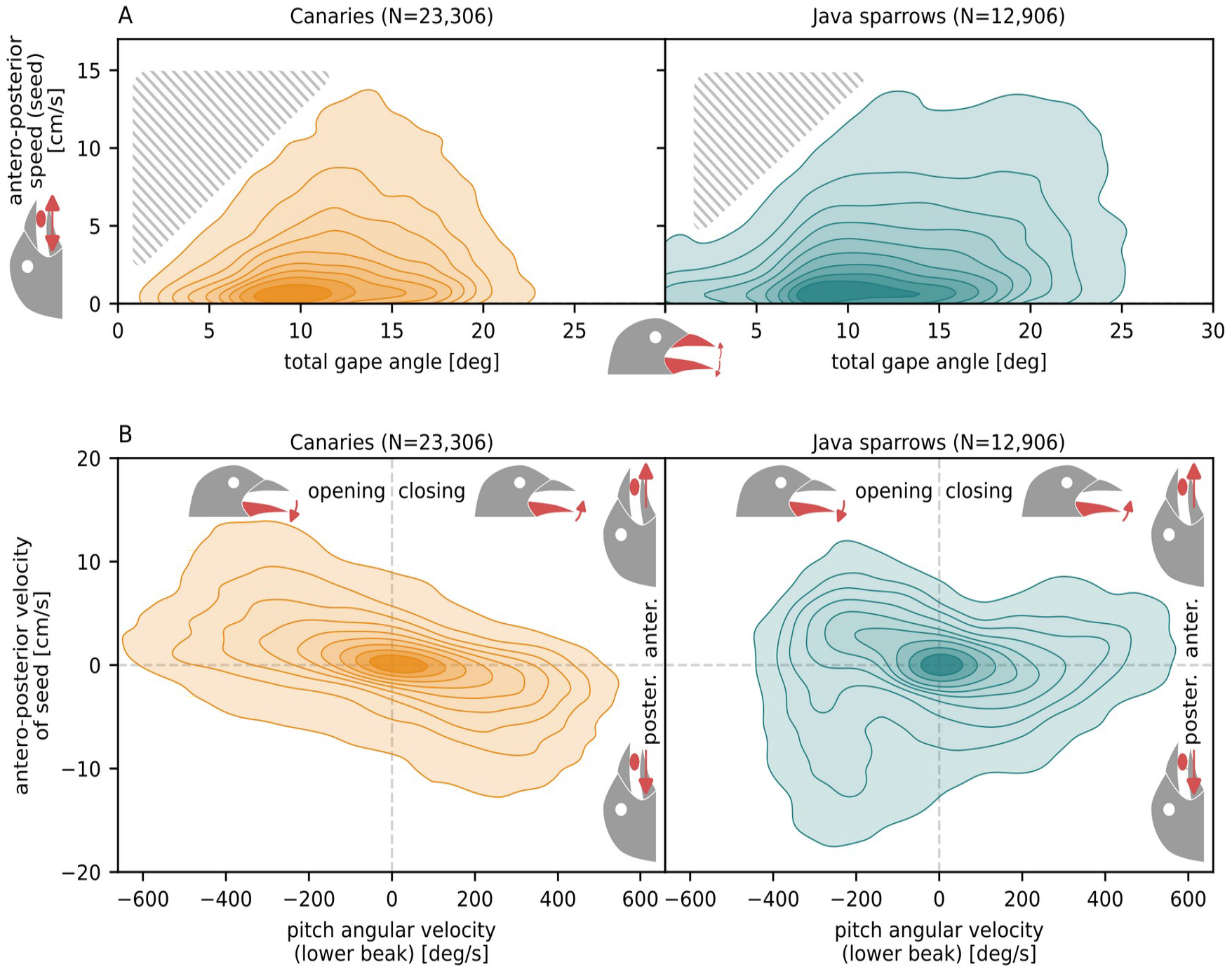
Coordination of beak movement and seed transport. Low speeds of seed movement at low gape angles (**A**, see gray hatched areas) indicate that the beak needs to release its grip to allow for fast seed movements. Canaries moved the seed mostly anteriorly during beak opening and posteriorly during beak closing **(B)**. Java sparrows showed a less specific pattern, with anterior and posterior seed transport being less strictly associated with either beak opening or closing. Plots show kernel density estimates of combinations of seed speed and gape angle (A) and seed velocity and mandible angular velocity **(B)**. More frequent combinations of values are indicated by darker colors. N denotes the number of frames included for calculation of kernel density estimates. All plots show the pooled data of two individuals per species.

Seed positioning involved cycles of beak opening and closing in which the beak alternately holds the seed for a brief moment and then widens the gape to allow for the seed to be moved by the tongue (Figure 7A). This implies that cycles of tongue movement occur at the same frequency as the beak cycles. The coordination of beak movement and seed transport differed between the species (Figure 7B). The relationship between seed translational velocity and lower beak angular velocity shows that canaries transported the seed posteriorly mostly during beak closing and anteriorly mostly during beak opening. In Java sparrows, we observed no such consistent association between direction of seed transport and beak opening or closing. We conclude that canaries use a more conserved coordination of beak movement and seed transport.

### Beak oscillation frequency during seed positioning

Given the many complex manipulations performed during feeding, a fast rate of beak opening and closing may be crucial for efficient seed processing. To quantify the frequency of beak oscillation, we calculated the main frequency of oscillation of the lower beak for each seed positioning event.

The main frequencies of lower beak movements used for seed positioning ranged from 6.6 to 24.7 Hz, with oscillation amplitudes between 2.3 and 10 degrees (Figure 8). When comparing the two species, we observed that canaries moved their beaks at faster rates than Java sparrows (mean frequency: 18.6±0.4 Hz vs. 13.5±0.4 Hz, 94%hdi_can-jav_=[4.0, 6.1]; see Figure 8 and Appendix 1—table 5A). Java sparrows showed a negative relationship between frequency and amplitude (regression slope: -1.2±0.4), indicating a limited ability for fast oscillations at high amplitudes (Figure 8B). In canaries, we observed a similar trend, but the relationship between frequency and amplitude was not significant (slope 94%hdi_can_=[-1.1, 0.4] includes zero, Appendix 1—table 5A). The slopes of the two species, however, were not significantly different from each other (Appendix 1—table 5A). In conclusion, songbirds use fast beak oscillations during seed positioning, which vary in speed among species and may be faster at lower amplitudes.

**Figure 8:**
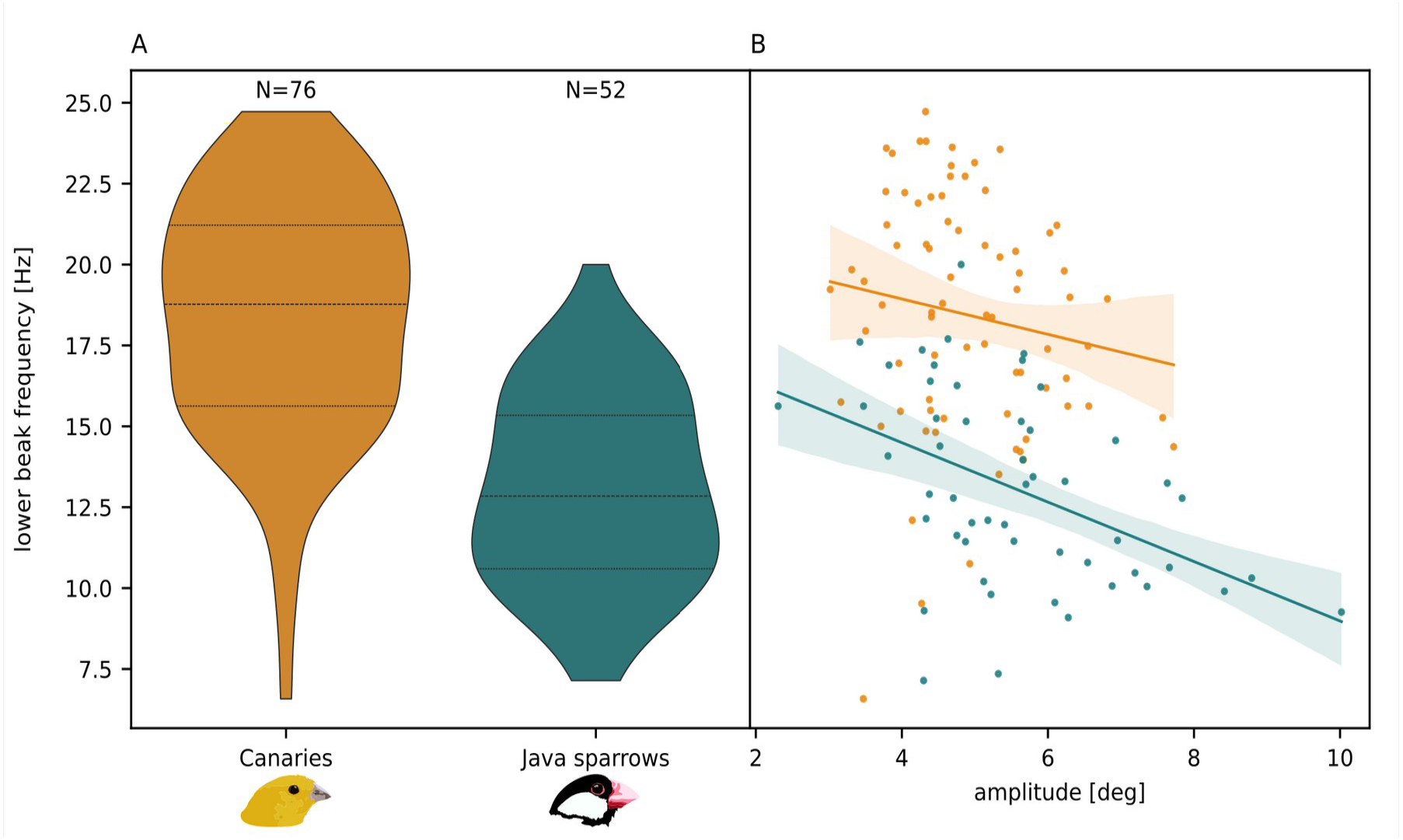
The frequency of lower beak oscillation differs between species **(A)** and may decrease with amplitude **(B)**. Plots show the lower beak pitch angle oscillation frequency during seed positioning. On average, the frequency is higher in canaries than in Java sparrows (18.6±0.4 Hz vs. 13.5±0.4 Hz, A). In Java sparrows, the frequency of oscillation decreases with increasing amplitude (B). This relationship is not significant in canaries. Dashed lines in A indicate quartiles of the distributions. Area in translucent shading in B indicates the 94% confidence interval of the regression. N denotes sample sizes, and the sample sizes per species are the same in B) as in A). Plots show the pooled data of two individuals per species. See Appendix 1—table 5A for results of the statistical analyses. **Source data 1.** Data of lower beak oscillation frequencies and amplitudes.

### Contraction and relaxation speed of jaw muscles

To assess whether the reason for slower oscillation rates in hard-biting Java sparrows lies in differences in intrinsic muscle contraction parameters, we conducted *in vitro* measurements of a major beak opener (*m. depressor mandibulae*, henceforth referred to as ‘depressor’) and a major beak closer muscle (*m. adductor mandibulae externus ventralis*, henceforth referred to as ‘adductor’). We electronically stimulated muscle preparations *in vitro* to produce isometric tetani and recorded force-time curves of the contractions (Figure 9 A+B). This allowed us to quantify half rise time (time after the start of muscle activation to reach 50% of maximum force) and half relaxation time (time after the start of muscle deactivation to drop to 50% of maximum force).

**Figure 9:**
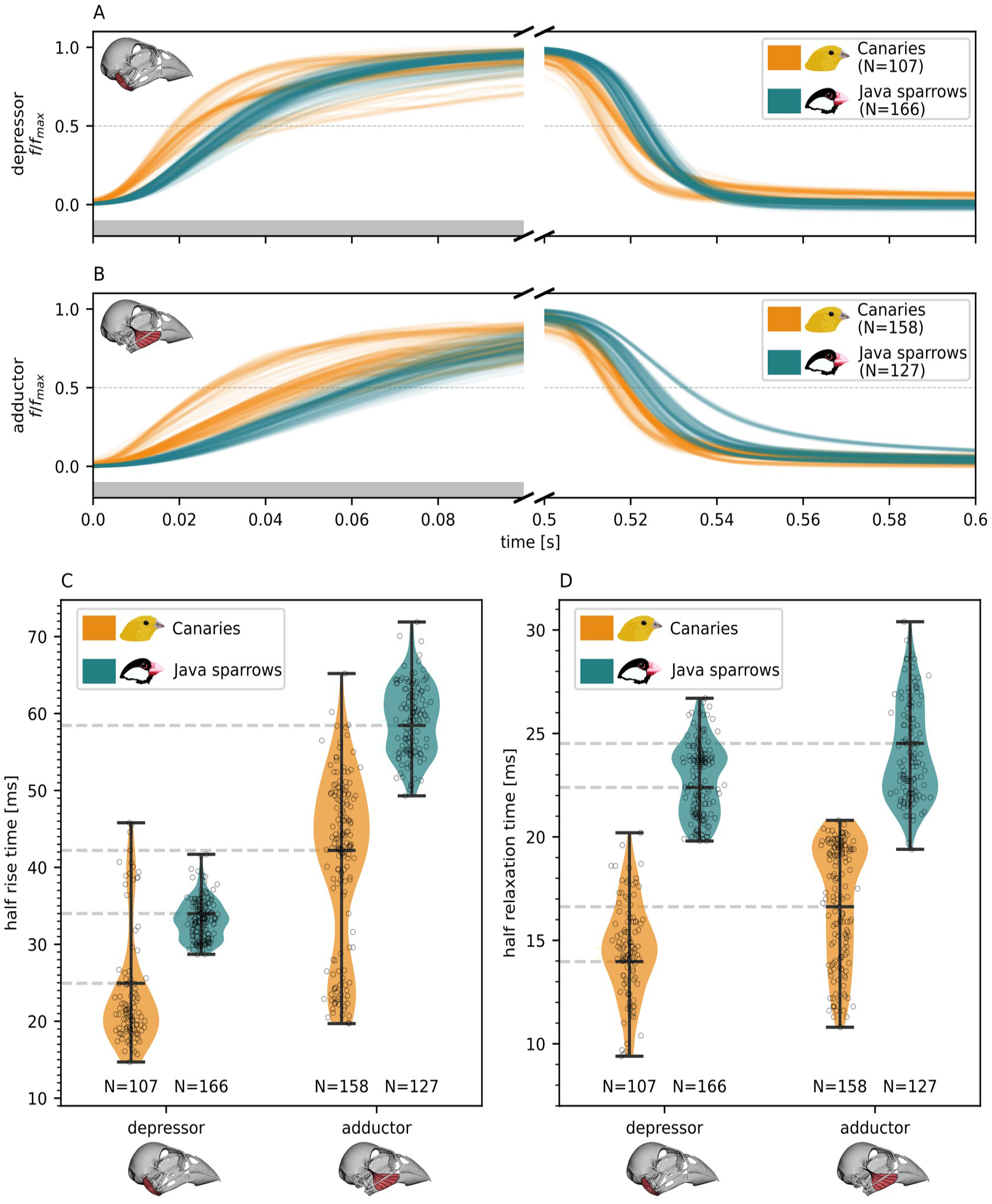
Contractile properties of adductor and depressor muscles. **A+B**) Force-time curves of isometric contraction of depressor and adductor, respectively. Force data are normalized by maximal force and show the active force only (passive force has been subtracted). The gray bar indicates the duration of the tetanus stimulation (start: 0 s, end: 0.5 s). Force curves in A and B show pooled data of four individuals per species and per muscle type. **C**) Time to reach 50% of maximal force during isometric muscle contraction under tetanus stimulation. **D**) Time to reach 50% of maximal force during muscle relaxation. N denotes sample sizes (number of tetani). Violin plots show the pooled data of four individuals per species and per muscle type. Gray hatched lines indicate mean values as obtained from statistical analyses (Appendix 1—table 3B). Source data 1. Half-rise times (C) and half-relaxation times (D) of jaw muscles.

Adductor muscles were significantly slower than depressor muscles in both contraction and relaxation, independent of the species (Figure 9 C+D, Appendix 1—table 5B). The differences between adductor and depressor were much higher for half rise times (69% and 72% for canaries and Java sparrows, respectively) than for half relaxation times (19% and 9% for canaries and Java sparrows, respectively). This indicates that relaxation time is more conserved across muscle types than rise time.

Both the adductor and the depressor muscles were slower in Java sparrows than in canaries (Figure 9, Appendix 1—table 5B). The depressor in Java sparrows was 37% slower during activation and 60% slower during relaxation compared to canaries. For the adductor, the differences were 39% and 48% for activation and relaxation, respectively.

Taken together, our results suggest that muscles optimized for high force production (adductor in contrast to depressor and muscles of Java sparrows in contrast to muscles of canaries) need more time to reach their maximal force after the start of activation and to relax after the start of deactivation.

## Discussion

Our data show that the tongue is an essential tool for efficient seed processing in small songbirds. Since finches do not use ballistic transport, they utilize their tongue as a unique tool to transport the seed relative to the beak and bring it into the desired position and orientation for a biting attempt (Zweers et al., 1994). We observed a high correlation of movement of the seed and tongue along the antero-posterior axis (Figure 2C), confirming a crucial role of tongue movements for seed transport within the beak. The tongue not only pushes the seed anteriorly but also pulls it posteriorly, sometimes quickly lifting the seed such that it loses contact with the beak. Furthermore, the birds utilize their tongue for the rotations necessary to optimize seed orientation and to stabilize the seed during biting attempts [Figure 4 C-E, cf. (Ziswiler, 1965)]. Morphological analyses of the tongues of canaries and other granivorous passerine birds (Başak et al., 2017; Dehkordi et al., 2010; Emura et al., 2010; Jackowiak et al., 2010; Toyoshima et al., 1992) revealed fine needle-like processes on the lateral sides of the lingual apex and backwards-pointing conical papillae on the dorsal posterior body of the tongue. It is possible that these morphological traits aid birds in the dexterous manipulation of food (Başak et al., 2017; Dehkordi et al., 2010; Jackowiak et al., 2010), though we currently lack empirical evidence to prove this. Taken together, the specialized morphological adaptations and the fast, coordinated movements of the tongue during feeding suggest granivorous songbirds rely heavily on their tongue as a major tool for seed processing.

Seed processing requires fast, precisely coordinated, and complex movements of the entire kinetic skull. The extensive usage of cranial kinesis in both species suggests a vital role of upper beak rotations for seed manipulation. Such rotations occur via the naso-frontal hinge (prokinesis) or via bending zones within the upper beak [rhynchokinesis and amphikinesis (Zusi, 1984)]. We could show that both tested species made extensive use of upper beak rotations during all phases of the feeding cycle. Previous research has shown that cranial kinesis is indeed an important feature for various avian feeding behaviors (Mielke & Van Wassenbergh, 2022; Bock, 1964, 1966; Zusi, 1984; Bout & Zweers, 2001; Estrella & Masero, 2007). It allows for wide beak gapes while keeping the depression of the lower jaw at moderate levels. This may enable the jaw adductor muscles to operate in more optimal length ranges, maintaining high force output even when handling large food items (Dickinson et al., 2022; Gidmark et al., 2013; Kaczmarek & Gidmark, 2020; Wilken et al., 2025). In granivorous songbirds, minimizing lower jaw depression may furthermore limit the risk of dropping the seed when keeping the head at a neutral position while looking out for predators (cf. Andries et al., 2023). Taken together, cranial kinesis facilitates both transmission of biting forces and careful manipulation of the seed.

The multiple degrees of freedom of the lower beak facilitate several aspects of the feeding process in granivorous songbirds. Longitudinal fore/aft movements (slicing) may be used to open closed-shelled seeds in other birds (Ziswiler, 1965) but did not play a major role in the feeding kinematics of the species tested here. Both species did, however, show considerable amounts of medio-lateral mandible rotations (yaw), which match the observations in other granivorous songbirds (Kear, 1962; Meij & Bout, 2006; Ziswiler, 1965). Mandible yaw helps to reduce husking time and the number of cracking attempts by optimizing the alignment of the lower beak with the seed (Meij & Bout, 2006). Multi-dimensional mandible movements seem particularly important during husk removal, as both species showed the widest range of yaw rotations during that phase. We suggest that lateral movements help to avoid or limit crushing of the seed while separating the husk from the inner kernel. During biting, yaw rotations may facilitate cracking of seeds in an upright or vertical-longitudinal orientation (cf. Figure 3B) by optimizing the alignment of the lower beak’s tomium with the edge of the seed where the two halves of the husk join and are cracked open most easily. During positioning, medio-lateral mandible movements play an essential role in seed rotation, as such types of complex manipulation require optimal arrangement of the food type with the tongue and the upper and lower jaws. Taken together, the complex manipulations necessary for efficient positioning, cracking, and dehusking of seeds would not be possible without multi-dimensional movements of the lower beak.

The partly independent movements of the upper and lower beaks may benefit from the absence of a postorbital ligament in some songbirds. In canaries, this absence allows them to flexibly adapt their beak movements to meet the specific requirements of the different phases of seed processing (Mielke & Van Wassenbergh, 2022), which contributes to efficient and careful seed processing and husk removal. Here, we showed that Java sparrows exhibit a stronger (negative) correlation of the upper and lower beaks than canaries, possibly caused by their strong postorbital ligament [‘coupled cranial kinesis,’ (cf. Bock, 1964)]. However, results from other studies question whether the postorbital ligament actually accounts for or is required for coupled cranial kinesis (Bout & Zweers, 2001; Hoese & Westneat, 1996; Lyons et al., 2023; Zusi, 1967). Further research is needed to reveal whether stronger coupling of beak movements in certain species is caused by morphological traits or whether it is a behavioral feature of their feeding technique.

Mechanoreception in the beak and tongue provides the basis for the fine sensory-motor control necessary for controlled seed manipulation (Gutiérrez-Ibáñez et al., 2026). The most common mechanoreceptors in birds are Herbst corpuscles, which occur in exceptionally high densities in the beaks of tactile foragers but also in other avian species (Ziolkowski et al., 2022). Herbst corpuscles are also present in the tongues of canaries (Başak et al., 2017) and other finches (Toyoshima et al., 1992). Previous studies have observed Herbst corpuscles in the dermis of the beak of Java sparrows, albeit at low densities compared to the lateral aspect of the upper beak (Genbrugge et al., 2012). However, the palate of Java sparrows is covered by an exceptionally thick epidermis (Genbrugge et al., 2012), which may impede the tactile sensation within the oral cavity necessary for fine-tuned sensory-motor control during seed handling. This may explain our observation that Java sparrows commonly crushed seeds during husk removal and showed a less consistent coordination of beak movement and seed transport. However, a quantitative and comparative analysis of mechanoreceptor abundance and thickness of the epidermis in hard- and weak-biting finches is needed to test whether a thick rhamphotheca may indeed impede sensory-motor control during seed handling.

The lower frequency of beak opening and closing in the harder-biting Java sparrows suggests a force-velocity trade-off may be at play. Their average frequency (13.5±0.4 Hz) was lower than that of canaries (18.6±0.4 Hz) and decreased more with increasing gape angles. These observations are in line with previous results on beak frequency in singing songbirds, which indicated that a force-velocity trade-off limits the speed of beak movement in hard-biting species (Demery et al., 2021; Derryberry et al., 2018; Friedman et al., 2019; García & Tubaro, 2018; Herrel et al., 2009). The underlying biomechanical mechanisms causing this trade-off may be differences in the cranial lever system, physiological properties of the jaw muscles driving beak movements, or a combination of these factors (Herrel et al., 2009).

Our results strongly suggest that the contractile properties of jaw muscles contribute to the force-velocity trade-off in beak movements of songbirds. We show that the muscles of a hard-biting species contract and relax at slower rates compared to those of a weak-biting species. We assume these contractile differences reflect variability in the relative proportions of muscle fiber types between the two species, where faster contraction and relaxation are assumed to be associated with increased amounts of fast-twitch fibers (Gronenberg et al., 1997; Jahromi & Atwood, 1969). Recent findings support this idea by showing that muscle fiber type strongly affects beak frequency output in simulations of mandible motions driven by Hill-type muscle models (Jorissen & van Wassenbergh, 2026). In general, our results confirm that *in vitro* contractile kinetics of muscles reflect their *in vivo* operating frequency (Gladman & Elemans, 2024).

The observed differences in beak kinematics, together with variations in beak morphology and bite force, reflect different dietary niche utilization. As harder-biting species have access to larger and harder food types, they need to consume fewer food items per day to meet their energy demands. In contrast, weak-biting songbirds like canaries are forced to meet their high energy demands by consuming high amounts of smaller and softer seeds. A small, fast, and agile beak, as in canaries, facilitates this feeding mode (Benkman & Pulliam, 1988; Smith, 1987). Previous studies have shown that small-billed finches may even be faster in processing small grains than large-billed species (Abbott et al., 1975). These findings suggest that not only the cranial morphology of granivorous songbirds (Felice et al., 2019) but also the kinematics of their feeding apparatus as a functional system is under strong selective pressures.

A potential limitation of our study is based on the fact that canaries and Java sparrows do not have the exact same skull size. This may have some limited impact on the results of our comparative analyses. The slightly smaller canaries have to handle a relatively larger seed at a shorter distance to the beak’s axes of rotation. To perform the same seed manipulations, they have to exert wider beak gapes than Java sparrows. Since gape size can affect bite performance (Dickinson et al., 2022), the feeding kinematics of the small canaries, as described here, may not be entirely representative of their manipulation of smaller-sized seeds. However, canaries show a strong preference for hemp seeds in food choice experiments (Andries et al., 2024). Therefore, our findings likely fall within the biologically meaningful range for canaries, although we note this may not be generalizable for other small passerines.

The larger relative seed size in canaries may explain the wider pitch and yaw angles that we observed, for example, during dehusking (Figure 5). However, we compared the observed rotation angles to those that canaries would theoretically need to perform to achieve the same amplitudes as Java sparrows during dehusking and found that canaries actually use 48% wider yaw and 21% wider pitch than would be necessary to compensate for their smaller beak size. This indicates that relative size differences have, if at all, only a minor effect on our results and do not compromise the conclusions drawn from the results.

Controlled object manipulation requires the coordinated, yet partially independent, movement of multiple elements, a principle exemplified by three-fingered robotic grippers (Kleinmann et al., 1996) and the primate hand (Elliott & Connolly, 1984). Our findings show that the avian feeding apparatus achieves comparable dexterous control through a three-element system comprising the upper beak, lower beak, and tongue. Unlike primate hands, which rely on a total of 27 degrees of freedom (ElKoura & Singh, 2003) and continuous multi-point contact (Elliott & Connolly, 1984), birds accomplish both forceful cracking and fine manipulation via coordinated, high-frequency repositioning cycles with a system of only six degrees of freedom (one for the upper beak, two for the lower beak, and three for the tongue). This demonstrates that dexterous manipulation can emerge from tightly integrated control of relatively few mechanical degrees of freedom.

## Materials and Methods

### Birds and housing

Domestic canaries (*Serinus canaria domestica*, Linnaeus 1758, Fife Fancy breed) and Java sparrows (*Padda oryzivora*, Linnaeus 1758, wild type) were housed in individual cages (60×50×40 cm) with food and water available *ad libitum*. The canaries originated from an outbred lab population of the Behavioral Ecology and Ecophysiology research group at the University of Antwerp, Belgium, and the Java sparrows were obtained from local breeders. All birds were adult males. The birds were given a standard seed mix, which contained hemp seeds, the seed type used for the feeding experiments. For the Java sparrows, we supplemented the seed mix with ∼25% paddy rice. Four birds (two individuals per species) were sampled for the XROMM workflow; 16 birds (8 individuals per species) were sampled for *in-vitro* muscle testing (see details below). The animal experiments performed for this study were approved by the Ethical Committee for Animal Testing of the University of Antwerp (ECD code: 2020-40).

### XROMM

We used marker-based X-ray reconstruction of moving morphology (XROMM) as originally described in Brainerd et al. (2010). XROMM allowed us to quantify the rotations of the upper and lower beaks in all dimensions relative to the skull. With the help of metal X-ray markers, we also described the movements of the tongue and the seed in 3D. Two individuals per species were taken through the XROMM procedure as described below.

#### Surgical marker implantation

Birds were put under general anesthesia for implantation of metal X-ray markers. Five hours before the start of the surgery, we moved the bird to the surgery room for acclimatization. One hour before the surgery, we induced fasting by removing food and water from the cage to ensure the bird went into surgery with an empty crop. The bird was weighed for calculation of the anesthetic dose. To facilitate marker implantation in the beak and tongue, we decided against mask induction with anesthetic gas. Instead, we induced anesthesia via intramuscular injection of a mix of ketamine (25 mg/kg) and medetomidine (2 mg/kg) in the superficial pectoral muscle, an established method for general anesthesia in zebra finches (Boumans et al., 2008). We used the toe pinch reflex to check for an adequate level of anesthesia. If needed, a 20% dose of the anesthetics was re-administered to reach or maintain the necessary level of anesthesia.

We implanted 0.35 mm spherical metal markers (alloy: Sn96.5Ag3Cu0.5) in the upper beak (4x), lower beak (4x), head (4x), and tongue (1x). For the beak markers, we used a micro hand drill to drill 0.35 mm holes in the rhamphotheca: two on the left and two on the right side of both the upper and the lower beak. Next, we pressed the metal markers into these holes and sealed them with tissue glue. Because of the hollow bones with an extremely thin cortex, it was not possible to insert the markers the same way into the skull bone. Instead, we placed the markers between the skull bone and scalp. To do so, we pressed each marker on the tip of a sterile hypodermic needle, inserted the needle with the marker under the scalp, and then gently pushed the marker out with a thin metal rod going through the hypodermic needle. If possible, the markers were fixed in place with tissue glue by dipping the needle tip with the marker into tissue glue before insertion under the scalp. The tongue marker was inserted into the middle of the tongue via a lateral injection using a hypodermic needle with a marker at the tip, which was again pushed out using a thin metal rod that went through the needle. To avoid accidentally sticking the tongue to the beak, we did not use tissue glue to seal the injection hole. In all but one bird, the tongue marker stayed in place for the whole duration of the X-ray recordings in the days after the surgery. The exception was a canary that had lost the tongue marker after the first recording day. For that bird, we only included trials of this first recording in the analyses.

During the surgery, we monitored the body temperature and breathing rate of the bird. Body temperature was measured with a rectal probe connected to a digital thermometer (ThermoWorks Therma Waterproof type T, resolution: 0.1°C). If possible, we measured the temperature continuously via the vent. If not possible, we measured the temperature in ∼5-min intervals on the abdominal skin. We helped maintain the body temperature using an adjustable heating pad under the bird and/or a heating lamp above the bird. The breathing rate was monitored continuously by the surgical assistant.

After finishing marker implantation, we waited for the first signs of recovery (first spontaneous movements of the bird) before administering 10 mg/kg atipamezole into the pectoral muscle to reverse the effect of medetomidine. Despite this antagonist, injected to aid in a smooth and rapid recovery (Ikebuchi et al., 2020; Memon et al., 2021), recovery took a long time, often with several hours passing before the birds were perching again. We conclude that further research is necessary to improve the anesthetic regime for general anesthesia induced via intramuscular injection in non-model songbird species.

During recovery, we closely monitored body temperature, body mass, and breathing rate, each measured at least once an hour (up to 4x per hour directly after the surgery). To ensure sufficient hydration in the warm environment, we drip-fed the birds with water using a pipette. We let the birds recover for 24 hours after surgery with food and water *ad libitum* before starting the fasting period for the feeding experiments.

#### X-ray recordings

For the X-ray experiments, we used hemp seeds (*Cannabis sativa*), a seed type that is typically used in experiments to study feeding in granivorous birds (Andries et al., 2023; Mielke & Van Wassenbergh, 2022; Van der Meij et al., 2004; Willson, 1972) and that was also part of the regular diet used to feed the birds outside of experiments. Hemp seeds have the shape of an elongate lens, with the shell made up of two halves joined at the outer edge of the lens. The hardness of hemp seeds is approximately 12 N (Van der Meij et al., 2004) but can be considerably higher at low humidity (Taheri-Garavand et al., 2012).

X-ray recordings of the birds with implanted metal markers were performed with the bi-planar 3D^2^YMOX setup of the Laboratory for Functional Morphology at the University of Antwerp. After at least 24 h recovery from the surgery, the birds fasted overnight (water available *ad libitum*) until the morning of the recording day. One hour before the start of the recordings, we moved the bird to the recording room for acclimatization. We used custom-built 3D calibration objects with metal markers to calibrate the system and recorded a fine mesh grid to allow for correction of image distortion later on. During the recordings, the birds were kept in a transparent plastic box with a feeder. The feeder with its short perch was placed such that the bird was positioned in the center of the focal volume while feeding. We offered hemp seeds that had a metal X-ray marker implanted in the center to make the seed visible in the recordings and to be able to reconstruct movements of the seed within the beak. Only one seed at a time was placed in the feeder (more were added if the bird was not motivated to feed). While the bird was feeding, we recorded videos of ∼6 s length at a frame rate of 500 frames per second and with an image resolution of 2048×2048 px. The length of recording trials was kept short to limit radiation exposure for the birds and prevent the system from overheating. The 6 s videos did not always capture the full feeding event, especially in canaries, who took longer to finish a seed compared to Java sparrows. But across all trials within a recording day, each feeding phase was captured at least four times for each individual. To later define the resting position of the upper and lower beaks (in closed position), we also recorded short sequences of the bird while it was not feeding. We also recorded videos synchronized to the biplanar X-ray recordings using a high-speed light camera (EOSENS-TS3 or FASTEC-IL5) using the same frame rate to record and later annotate the different phases of seed processing. To limit radiation exposure, we recorded no more than 20 trials per recording day. Each bird was recorded on two days, with one to three days in between without X-ray exposure.

#### CT scanning and segmentation

Upon completion of the last recording day, the birds were euthanized via an intracoelomic injection of 0.05 ml sodium pentobarbital (200 mg/ml). Directly after, they were CT-scanned at the DynXLab at the University of Antwerp (DynXLab: Center for 4D quantitative x-ray imaging and analysis) with 15 μm (3 birds) or 18 μm (1 bird) voxel size. The scans were imported into the Dragonfly software, version 2022.2 (Dragonfly 2022.2 for Windows, 2022), to segment the rigid bodies of interest (braincase and the upper and lower beaks, including rhamphotheca) and to extract the 3D positions of the implanted metal markers. The segmented regions were transformed to surface meshes and exported.

#### Marker tracking

Marker tracking was conducted in XMALab version 2.1.0 (Knörlein et al., 2016). If necessary, the X-ray videos were improved via flat-field correction beforehand, using empty background recordings taken during the experiments. Using the built-in functions for calibration and undistortion in the XMALab software, we prepared an ‘.xma’ file for each recording day. The trials were imported into the dataset, and the positions of the 14 X-ray markers (4x head, 4x upper beak, 4x lower beak, 1x tongue, 1x seed) were tracked automatically via dot detection. Via evaluation of reprojection errors of each individual marker and of marker-to-marker distances of markers that were part of the same rigid body, incorrectly tracked frames were identified and then manually corrected. The tracking precision (mean standard deviation of inter-marker distances of markers from the same rigid body) was 0.05 mm. For further processing of the data in Autodesk Maya (see below), we exported camera positions, undistorted trial images, 3D positions of the markers, and transformation data of the rigid bodies. During export, 3D positions and rigid body transformations were low-pass filtered with a threshold of 60 Hz.

#### XROMM animation and data export

In Autodesk Maya 2023.3 (Autodesk, INC., 2023), we imported the 3D models of the braincase, upper beak, and lower beak and animated the meshes with the rigid body transformation data exported from XMALab. At the beginning of each video, we added one frame with the meshes arranged as a closed beak, retrieved from the recordings of non-feeding trials. This closed beak arrangement was used to define and position the anatomical coordinate system (ACS) and the joint coordinate systems (JCS) as follows (cf. Figure 1D):

First, to make sure that all coordinate systems are placed within the plane of symmetry of the skull, we defined the median plane as a surface that is defined by the following three points: 1) the tip of the beak in closed position, 2) the central point of the dorsal edge of the foramen magnum, and 3) the central point of the naso-frontal hinge. Both the ACS and the two JCSs were positioned such that the x-axes and y-axes lie within this median plane; the x-axes run through the tip of the beak, and the y-axes point dorsally. All z-axes were oriented to the right. The origin of the ACS was placed at the central point of the dorsal edge of the foramen magnum. The origin of the JCS for the upper beak was positioned in the center of the naso-frontal hinge. The origin of the JCS for the lower beak was positioned such that the z-axis ran through the centers of the left and right quadrato-mandibular joints (Figure 1D). The JCSs of the upper and lower beaks were defined with the braincase as the proximal element. This way, kinematic data quantify the beak movements relative to a virtually fixed cranium.

For both JCSs, we exported the rotation angles around the z-axis (pitch), and for the lower beak also the angles of rotation around the y-axis (yaw). Furthermore, we exported the 3D positions of the seed marker and the tongue marker relative to the ACS. These raw data were deposited on Dryad (link pending).

We did not include translations of the lower jaw in the data analysis. Such translations are theoretically possible due to the suspension of the mandible via the quadrate bones in avian skulls. In our data, the average standard deviation of lower beak translations stayed below 0.35 mm. This contribution to overall beak kinematics was insignificant compared to rotational movements around the quadrato-mandibular joints (cf. Appendix 1—figure 1 and Appendix 1— figure 2).

### Processing of XROMM data

First, we annotated the different phase types (pick-up, positioning, biting, dehusking, and swallowing) in each frame of each video to be able to filter for specific phase types later on in the analyses. To annotate phase types, we used the light camera videos that we recorded in sync with the X-ray videos. For this study, we focused on the phase types ‘positioning,’ ‘biting,’ and ‘dehusking.’ Seed positioning was defined as sequences during which a bird oscillated the beak and moved the (still intact) seed within the beak with the tongue. A biting phase was defined as the frames in which the seed is fixed between the upper and lower beaks and vertical forces are applied. The start of a dehusking phase was defined as the moment in which the first crack in the husk appears. The end of dehusking was defined as the moment in which the last piece of husk fell out of the beak. In Java sparrows, this was difficult to detect in some trials, as they often split the husk into many fragments. In this case, we defined the end of dehusking as the moment in which the bird starts swallowing, i.e., starts to move the seed towards the esophagus by antero-posterior movements of the tongue (as visible in the X-ray recordings thanks to the metal markers in the tongue and the seed).

#### Maximum gapes

To analyze the maximal beak opening during the different feeding phases, we extracted local maxima in total gape angle over time. To do so, we first calculated the total gape angle as α_total_=α_upper_-α_lower_, with α being the pitch angle (α_lower_<0 for depressed lower beak). Next, we extracted local maxima in total gape within each event of positioning, biting, and dehusking using the `scipy.signal.find_peaks()` function in Python. For each frame in which a local maximum was found, we extracted the total gape itself as well as the pitch angles of the upper and lower beaks (see Figure 5 A+B). The data are available in Figure 5–Source Data 1.

#### Frequency of beak oscillation

To calculate the frequency of beak movement, we extracted the main oscillation frequency from the pitch angle data of the lower beak per positioning event. We chose that phase type because during positioning, the beak moves with higher amplitudes and at more regular rates compared to the other phase types. If a positioning event was not tracked entirely (e.g., because of markers being untrackable because of motion blur during fast head movements of the bird), we processed the fully tracked sequences of frames within an event separately. Only sequences of at least 50 frames in length were processed.

As preparation, we first conducted an empirical mode decomposition [EMD, (Zeiler et al., 2010)] on each sequence of interest. This allowed us to subtract the low-frequency local trend from the signal, which simplified detection of the actual oscillation frequency that we were interested in. Furthermore, it allowed us to calculate a reasonable proxy for oscillation amplitude (see step 2 below). The EMD included three steps: 1) We detected local maxima and minima in the lower beak’s pitch angle over time using the `scipy.signal.find_peaks()` function in Python. We proceeded with the sequence only if at least three maxima and three minima were found. 2) We interpolated the range between neighboring maxima and neighboring minima using the `scipy.interpolate.pchip_interpolate()` function to define an upper and lower ‘envelope’ for the signal. The mean distance between upper and lower envelopes was defined as a proxy for oscillation amplitude for that sequence. 3) We calculated the local trend as the mean value between the upper and lower envelopes and subtracted this low-frequency component from the data to facilitate detection of the actual feeding-related oscillation frequency.

Next, we extracted the main oscillation frequency from the processed data of each sequence. As preparation, we first filtered the data with high-pass and low-pass filters with thresholds of 6 Hz and 60 Hz, respectively, using the Butterworth filter [`scipy.signal.butter()`], applied forward and backward to the signal [`scipy.signal.filtfilt()`] to avoid introducing phase shift. Then we used Welch’s method with `scipy.signal.welch()` to calculate the power spectral density of the signal. This allowed us to extract the main frequency (the frequency with the highest power) from the signal. If there was more than one main frequency (more than one local maximum in the power spectrum), we discarded the sequence if the difference in power between the two strongest frequencies in the signal was smaller than 0.1. That is, if there was no clear candidate for the most prominent frequency in the signal. The data are available in Figure 8–Source Data 1.

#### Yaw angles of the lower beak

To analyze the lateral rotations of the lower beak, we extracted the yaw angles from the JCS for the three phase types of interest (positioning, biting, and dehusking, Figure 5C). To achieve the best consistency among trials, we centered the yaw angle data of each trial around the trial mean so that each trial mean is fixed at zero degrees.

To quantify the extent of lateral excursions of the lower beak, we extracted the standard deviation of yaw angles from each event of the phase types of interest. High standard deviations indicate wide lateral excursions of the lower beak. The resulting distributions of standard deviations could then be compared between the species (Figure 5D). The data are available in Figure 5–Source Data 2.

#### Correlation of upper and lower beak

To analyze to what extent the opening and closing of the upper and lower beaks are coupled, we analyzed the relationship between their pitch angular velocities (Figure 6). Angular velocities were extracted from the pitch angle data using the `numpy.gradient()` function in Python. The results provide information about the direction of rotation (positive and negative velocities indicating elevation and depression, respectively) as well as the magnitude of change (high absolute values indicating fast change). For each phase type of interest (positioning, biting, and dehusking), we extracted frame-wise pitch angular velocities for the upper and lower beaks (available in the Dryad dataset—link pending). If, for a given frame, the angular velocities of the upper and lower beaks are both positive, then both rotate upwards at that moment. If both are negative, they rotate downward together. If one value is positive and the other negative, then they rotate in opposite directions. The more balanced the distribution of angular velocity combinations (i.e., the more the rotations deviate from the standard opposing directions), the more independent the rotation of the upper and lower beaks. See below (‘Quantification of correlation’) for details on how we quantified the correlation of upper and lower beak pitch angle time series.

#### Coordination of beak movement and seed transport

To analyze the coordination of beak movement and seed transport, we calculated the framewise pitch angular velocities of the lower beak and the translational velocities of the seed marker using the `numpy.gradient()` function in Python (similar to analysis of angular velocity of the upper and lower beaks; see above). These data are included in the Dryad dataset (link pending). For the seed, we considered only translations along the antero-posterior (x-) axis of the ACS, because most translations occur in this direction during positioning. This analysis allowed us to explore whether the seed is transported in a specific direction during beak opening or closing (Figure 7), which would indicate that the bird coordinates beak movements and seed transport.

#### Function of the tongue for seed transport and stabilization

To analyze the relationship between movements of the tongue and seed, we calculated the framewise velocities of the tongue and seed markers using the `numpy.gradient()` function in Python (similar to analysis of angular velocity of the upper and lower beaks; see above). These data are included in the Dryad dataset (link pending). Velocities were extracted from the translation data of the seed and tongue markers along the three dimensions defined by the ACS (Figures 2 + 4). If, for a given axis, velocities of the seed and tongue markers are both positive or both negative, then the tongue and seed move in the same direction along this axis, indicating a seed transport function of the tongue.

Since integrity of the seed is not a given anymore during the dehusking phase, and the seed husk rather than the kernel containing the marker is moved by beak and tongue, we did not include the dehusking phase in this analysis. To quantify the correlation between translations of the seed and tongue markers, we implemented a permutation test (see below).

### Quantification of correlation

To quantify the correlation of upper and lower beak pitch rotations (Figure 6) and of the three-dimensional translations of the seed and the tongue (Figures 2C-E and 4C-E), we implemented a permutation test using a modification of the concepts described by Yuan & Shou (2024, 2026). Strong temporal autocorrelation in time series data complicates the interpretation of correlations because parameters can easily yield strong correlation signals that do not necessarily reflect the true relationship of the data (Yuan & Shou, 2024; Yule, 1926). Permutation tests help to assess the correlation of time-series data more reliably by comparing the observed correlation with correlations obtained when one of the time series is replaced by surrogate data (Lancaster et al., 2018; Yuan & Shou, 2024, 2026). Such a surrogate can be, e.g., data from another trial (between-trial permutation) or a time-shifted sequence from within the same trial (within-trial permutation). For our particular experimental design (large number of trials of different lengths), the characteristics of our data (often oscillatory), and our specific research question (quantify the correlation for sub-data sets of certain species-phasetype combinations), it was reasonable to apply a test that combines within-trial and between-trial permutations. The procedure, described in detail below and visualized in Appendix 1—Figure 7, was implemented in Python and is provided as Supplementary file 2. The data file processed by the Python script is available in the Dryad dataset (link pending).

First, we extracted data slices from the chunks of kinematic data (step 1 in Appendix 1—Figure 7). Here, a ‘chunk’ is a continuous sequence of tracked frames of a certain phasetype in a trial. A ‘slice’ is a sequence of a defined number of frames extracted from a chunk. We used a slice length of 65 frames (130 ms), which covers a minimum of one full beak opening-closing oscillation cycle for most of the frequencies measured (cf. Figure 8A). Chunks of less than 65 frames in length were not included in this analysis. Slices are a collection of shifted time series from a chunk with an overlap of slice length minus 1 frame (hence, 64 frames). That means, the first slice of a chunk covers the frames 1-65, the second slice frames 2-66, etc. We chose a step size of 1 frame by default, as this extracts all possible slices from the data and does not require the user to select and justify a step-size parameter, which minimizes potential user-introduced bias in the analysis.

Next, for each parameter pair of interest (either upper and lower beak pitch rotations or seed and tongue translations), we calculated the cross-correlation matrix of all the slices extracted for each individual and phasetype (step 2 in Appendix 1—Figure 7). Each value in this matrix lies between -1 and 1, with values around zero indicating no correlation and values close to -1 and 1 indicating strong negative and positive correlation, respectively. In this matrix, the main diagonal represents the set of correlation values of the observed, non-shifted time series slices. The off-diagonal values represent the correlation values of permuted slice pairs.

To assess the correlation of the observed data, we compared the distributions of correlation values between the observed data and permuted data (steps 3 and 4 in Appendix 1—Figure 7). To do so, we calculated the median and the skewness of each of the distributions. The median represents the 50th percentile of the correlation and has a value close to zero in case there is no correlation in the dataset. The skewness was quantified via fitting a Beta distribution to the correlation values (after transforming the data from range [-1,1] to range [0,1]). With α and β being the shape parameters of the Beta distribution, skewness *sk* is defined as:

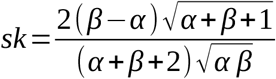

This parameter quantifies the degree of asymmetry of the distribution, indicating how much the values in the dataset lean towards the left (*sk*>0) or right (*sk*<0). The correlation values of permuted slices represent a randomized control and are expected to yield an approximately symmetrical distribution with median and skew values around zero (which we observed). The stronger the observed data deviates from the permuted data, the more likely it is that the measured correlation signal is not due to chance.

### Muscle physiology experiments

For the recording of *in vitro* force curves of beak opener and beak closer muscles, we used an Aurora Scientific Dual-mode Lever System (300C series, Aurora Scientific, ON, Canada) with a 300C muscle lever, which allows for force recordings with resolutions of 0.3 mN. Stimulation protocols were executed via the Dynamic Muscle Control software (DMC version 5.3, Aurora Scientific). This experiment was conducted with the major beak opener muscle (*m. depressor mandibulae*) and a major beak closer muscle (*m. adductor mandibulae externus ventralis*; see Figure 1E). For each muscle type in each species, four individuals were sampled. All birds were adult males.

#### Muscle dissection and mounting

Birds were euthanized via an intracoelomic injection of 0.05 ml sodium pentobarbital (300 mg/ml), approx. 5 mm cranial to the vent, below the keel. Death was confirmed by the absence of breathing and by testing both palpebral and toe pinch reflexes. After confirmation of death, the head was cut off and the skin removed on the left side of the skull to reveal the underlying jaw muscles. Next, the head was immersed in avian dissection buffer [150 mM NaCl, 2.5 mM KCl, 0.5 mM CaCl_2_, 1 mM NaH_2_PO_4_, 6.5 mM MgSO_4_, 10 mM HEPES, 12 mM glucose, adjusted to pH 7.4 with a 1 M Trizma base solution, cf. (Adam et al., 2021)] that was chilled with a bath of ice water and continuously oxygenated. Under a microscope (Leica M165 C), fascial and fat tissue was removed, and the muscle of interest was dissected out, leaving pieces of bone on each side of the muscle. After removing any remaining unwanted tissue from the muscle, we inserted suture loops in both pieces of bone attached to the muscle using thin metal wires. For the adductor muscle, we had to insert the suture through the muscle and the *fenestra mandibulae caudalis* because the muscle covers the entire lateral side of this part of the lower jaw. In doing so, we made sure not to rupture any muscle fascicles but rather carefully insert the suture material between the fascicles. Next, we transferred the muscle into the recording chamber, filled with warmed (37.2±0.9°C) avian recording buffer [150 mM NaCl, 2.5 mM KCl, 4 mM CaCl _2_, 1 mM NaH_2_PO_4_, 1 mM MgSO_4_, 10 mM HEPES, 12 mM glucose, adjusted to pH 7.4 with a 1 M Trizma base solution, cf. (Adam et al., 2021)], where we attached the muscle with the suture loops to a static hook on one side and to the force lever on the other side. After carefully adjusting the distance between the hooks until the muscle was slightly stretched (until we recorded a passive force between 10 and 20 mN), we let the muscle rest until the passive force reached a steady state. During the entire time the muscle was in the recording chamber, we provided continuous oxygenation and constant flushing with fresh, warmed recording buffer.

#### Isometric force recordings

To record force curves of the mounted muscle, we induced contraction with tetanus stimulations. Each protocol started with a 0.2 s initial delay to record the passive force, followed by a tetanus stimulation for a duration of 0.5 s with pulses of 0.2 ms length. The force was recorded with a sampling rate of 10,000 Hz.

First, we optimized the frequency of the tetanus stimulation and the length of the muscle. The purpose of the optimization was to obtain force curves with a maximum peak force (optimal signal-to-noise ratio) and with a plateau phase. To optimize the frequency, we first stimulated the muscle with different frequencies, increasing at intervals of 50 Hz. The frequency *f_o1_* that produced the best results was then further optimized by comparing force curves for *f_o1_*-25 Hz, *f_o1_*, and *f_o1_*+25 Hz. The best of these three frequencies was chosen as the optimal frequency, *f_o2_*. To optimize muscle length, we stretched the muscle at intervals of 0.2 mm and measured the force response to stimulation with *f_o2_* for each length. After each length increment, we waited for the passive force to reach steady state before we stimulated the muscle, which could take up to ∼15 min. The optimal length, *l_o_*, was defined as the length that did not show an improved force curve compared to the previous tested length. After obtaining *l_o_*, we repeated the frequency optimization as described above to obtain the final optimal frequency *f_o_*, which lay between 250 Hz and 550 Hz depending on species and muscle type.

Next, we recorded multiple force curves at *l_o_* and with *f_o_* to obtain half-rise and half-relaxation times. Between stimulations, we let the muscle rest for a minimum of three minutes. We repeated these measurements as long as the muscle provided a force response with a reasonable signal-to-noise ratio. For the analysis, we only included recordings with an active force (measured force minus passive force) of at least 6 mN. All raw force data were exported unfiltered and further processed in Python.

To reduce the noise, we filtered the raw force data with a low-pass filter (threshold 25 Hz) using the Butterworth function `scipy.signal.butter()`, applied forward and backward to the signal [`scipy.signal.filtfilt()`] to avoid introducing phase shift. To obtain the active force of the muscle, we subtracted the passive force (the mean force during the 0.2 s before the start of stimulation) from each data point. To obtain half-rise time *t_ri_*and half-relaxation time *t_re_* (Figure 9 C+D), we first measured the peak force *f_max_* in each active force curve and the latency times of muscle activation (*t_la_*) and deactivation (*t_ld_*). The latency time of muscle activation, *t_la_*, was defined as the time between the start of the muscle stimulation and the moment at which the filtered force curve crossed the threshold *f_at_=f_am_+3⋅f_asd_*, with *f_am_* and *f_asd_* being the mean and standard deviation of the unfiltered force data during the 50 ms before the start of stimulation. Accordingly, the latency time of muscle deactivation, *t_ld_*, was defined as the time between the end of the muscle stimulation and the moment at which the filtered force curve crossed the threshold *f_dt_=f_dm_−3⋅f_dsd_*, with *f_dm_* and *f_dsd_* being the mean and standard deviation of the unfiltered force data during the 50 ms before the end of stimulation. We then measured half-rise time *t_ri_* as the time between *t_la_* and the moment at which the filtered force curve reached *1/2f_max_* during force rise. Accordingly, we measured half-relaxation time *t_re_* as the time between *t_ld_* and the moment at which the filtered force curve dropped to *1/2f_max_* during relaxation. Half-rise and half-relaxation times are available in Figure 9–Source Data 1.

### Statistical analyses

#### General explanations

We applied statistical modeling analyses based on Bayesian sampling, conducted in Python (version 3.12.8, http://www.python.org) using the library “Bambi” [version 0.15.0, (Capretto et al., 2022)]. “Bambi” is a Bayesian model-building interface that is built on top of the probabilistic programming framework “PyMC” [version 5.20.0, (Salvatier et al., 2016)], which uses the NUTS sampler as the default sampling method. We ran relatively simple models, all with the ‘Gaussian’ model family and ‘identity’ link function. We sampled four chains, retaining 30,000 draws of each chain (which already exclude the default 1,000 iterations run to tune in the sampling algorithm). This resulted in posterior sample traces of length n=120,000. Model diagnostics (e.g., effective sample size, Gelman-Rubin statistics) were inspected but gave no signs of sampler issues.

Using the model, we asserted how much canaries (*can*) and Java sparrows (*jav*) differ regarding the various measures of jaw angular kinematics and muscle dynamics and how likely it is on the background of parameter variability that an observed difference is meaningful. For the muscle contraction parameters, we additionally quantified the differences between muscle types within each species following the same approach as explained below for inter-species differences.

The quantitative modeling approach was chosen over conventional hypothesis tests because of the sample size structure within our data: we obtained a relatively high number of observations for each individual, but from a low number of individuals that further belong to either of the two species of interest. This is a typical situation in XROMM and muscle physiology experiments. Sample size on the individual level was limited to four (two per tested species) for the XROMM experiments and 16 for the muscle measurements (four per combination of species and muscle type). Overall, this is too low for ‘individual’ to be included as a nested covariate in the model function together with ‘species’ (as no individual belongs to two species, these parameters must be nested). To still incorporate ‘individual’ as a model parameter and get species differences, we processed the data as follows (see Appendix 1—Figure 8 for a visualization of the workflow).

First, we generated posterior sample traces of the model parameter for each of the individuals (two canaries and two Java sparrows for the XROMM data, four canaries and four Java sparrows for the muscle data per muscle type). Posterior sample traces are the best attempts of the model to fit the model parameters to match the observed experimental data. The basic model functions (used consistently, unless stated otherwise below) to infer a parameter *θ* was *θ_i_=α_i_+ε*, or in short *θ∼individual*. Note that, because residual model variation *ε* is conventionally incorporated, parameter *θ* should be interpreted as the individual mean, and its sample variability as the variability of the mean. Each individual gets an independent sample trace of iterations, but for the same number of iterations (index *t*) and processed in parallel. For example, the posterior sample of parameter *θ* for individual *i* has the form *θ_i_=*[*θ_1,i_ θ_2,i_ … θ_n,i_*]=[*θ*]*^n^*, stored in the form of a column vector. These traces are concatenated to a table {*θ_t,i_*}, which shows the four individuals in columns and posterior sample traces in rows (Appendix 1—Figure 8). Such trace tables are the conventional outcome of probabilistic models, usually summarized immediately to retrieve the mean, standard deviation, and hdi (highest density interval, i.e., the smallest possible interval that includes a given fraction of the probability density). In this case, we further processed the raw table to get from ‘individual’ traces to conclusions on the ‘species’ level.

This was achieved by computing species averages per iteration *t* from the posterior samples, adding one new column *θ_s_* per species *s* with 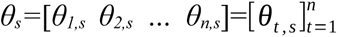. Species summary statistics (as presented in the tables in Appendix 1) were calculated for *θ_can_* and *θ_jav_* by computing the mean, standard deviation, and the 94%hdi over the rows. Finally, we calculated the sample-wise differences between the two species as 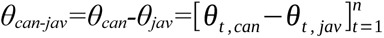 The summary statistics of *θ_can-jav_* (mean, standard deviation, and the 94%hdi over all samples) are presented in the tables in Appendix 1. If the 94%hdi of *θ_can-jav_* did not include zero, we concluded that there was a significant difference between species.

To reiterate, the parameter differences we calculated are the outcome of a probabilistic model that attempts to fit model parameters as a normal distribution to the observed data. It covers the variability in individual mean on parameter posterior sample traces and residual variability in the actual observations on model residual variability. Individuals per species are averaged *ex post*, and the calculated species difference traces cover the range of plausible true parameter values, given the experimental data. If the 94%hdi of the species difference does not cover zero, we interpret that as a meaningful difference. Python scripts used for the statistical analyses are provided in Supplementary file 1.

#### Maximal pitch angle

From the pitch angle data of the JCSs, we derived local maxima in total gape angle and the corresponding pitch angles of the upper and lower beaks, as visualized in Figure 5 A+B. For each of these three variables, we fitted the model "`angle∼individual`" to generate posterior samples approximating the mean angles for each of the four individuals. From there, we followed the approach as described above to test for significant differences between the species.

#### Oscillation frequency of the lower beak as function of amplitude

For the frequency of beak oscillation, we were interested in both the effect of species and the effect of the oscillation amplitude on the data. To generate posterior samples approximating the mean frequency per individual while accounting for potential effects of ‘amplitude,’ we included the latter as an additional main effect ("`frequency∼individual*amplitude`"). Here, the `*` operator denotes full interaction; "`individual*amplitude`" is shorthand for "`individual+amplitude+individual:amplitude`" (Capretto et al., 2022). Finally, we again followed the approach as described above to test for significant differences between the species.

#### Contractile properties of jaw muscles

From the force-time curves of *in vitro* muscle stimulation, we extracted half-rise and half-relaxation times of adductor and depressor muscles of both species. The statistical analysis addressed two questions: 1) Is there a significant difference between the two species in the contractile properties of the same muscle type? 2) Within each species, is there a significant difference in contractile properties between the two muscle types? For both half-rise time and half-relaxation time, we fitted the model "`time∼individual`" to generate posterior samples per individual and then followed the same approach as described above to test for significant differences between the species or between the muscle types.

#### Medio-lateral movement of the lower beak (yaw angles)

To quantify the width of the range of lower beak yaw rotations, we calculated the standard deviations of used yaw angles for each event of the phase types positioning, biting, and dehusking. Then, we tested whether the standard deviation differs between the species for each phase type. To do so, we fitted the model "`stdev∼individual`" to generate posterior samples of the mean standard deviation per individual and subsequently followed the approach as described above to test for significant differences between the species.

## Acknowledgments

We thank Wendt Müller for providing the canaries, Peter Scheys for help with animal care and housing, Gunther Vrolix for providing anesthetics for bird surgeries, Joaquim Sanctorum for the great CT-scans, Nathalie Van Houtte for providing chemicals, lab material, and technical advice, and Melanie Brauckhoff for an introduction to avian jaw muscle dissection techniques.

## Additional information

### Author contributions

Maja Mielke, Conceptualization, Methodology, Software, Formal analysis, Investigation, Visualization, Writing— original draft, Writing**—**review & editing, Funding acquisition; Falk Mielke, Methodology, Software, Formal analysis, Investigation, Writing**—**review & editing; Nicholas W. Gladman, Methodology, Writing**—**review & editing; Dan A. Tatulescu, Formal analysis, Visualization, Writing**—**review & editing; Anthony Herrel, Resources, Writing**—**review & editing, Funding acquisition; Coen P. H. Elemans, Methodology, Writing**—**review & editing; Sam Van Wassenbergh, Conceptualization, Methodology, Investigation, Supervision, Writing

## Additional files

### Videos

**Video 1.** Examples of the different types of seed rotation in canaries and Java sparrows.

**Video 2.** Example sequences of the XROMM animations of a canary and a Java sparrow. Positions of seed and tongue markers are indicated by orange and pink spheres, respectively.

**Video 3.** Example sequences of husk removal in a canary and a Java sparrow.

### Source Data files

**Figure 5-Source data 1.** Data of maximal total gape angles and associated pitch angles of upper and lower beaks.

**Figure 5-Source data 2.** Standard deviations of lower beak yaw rotations.

**Figure 8-Source data 1.** Data of lower beak oscillation frequencies and amplitudes.

**Figure 9-Source data 1.** Half-rise times and half-relaxation times of jaw muscles.

### Supplementary files

MDAR checklist

Supplementary file 1. Python script used for statistical analysis.

Supplementary file 2. Python script used for analyses of correlation of time series.

### Data availability

The raw XROMM data have been uploaded as a Dryad dataset which is currently kept private for peer review. Any additional data are provided as source data files. The Python scripts used for statistical analysis are available in the Supplementary files.

## Appendix 1

**Appendix 1—figure 1.**
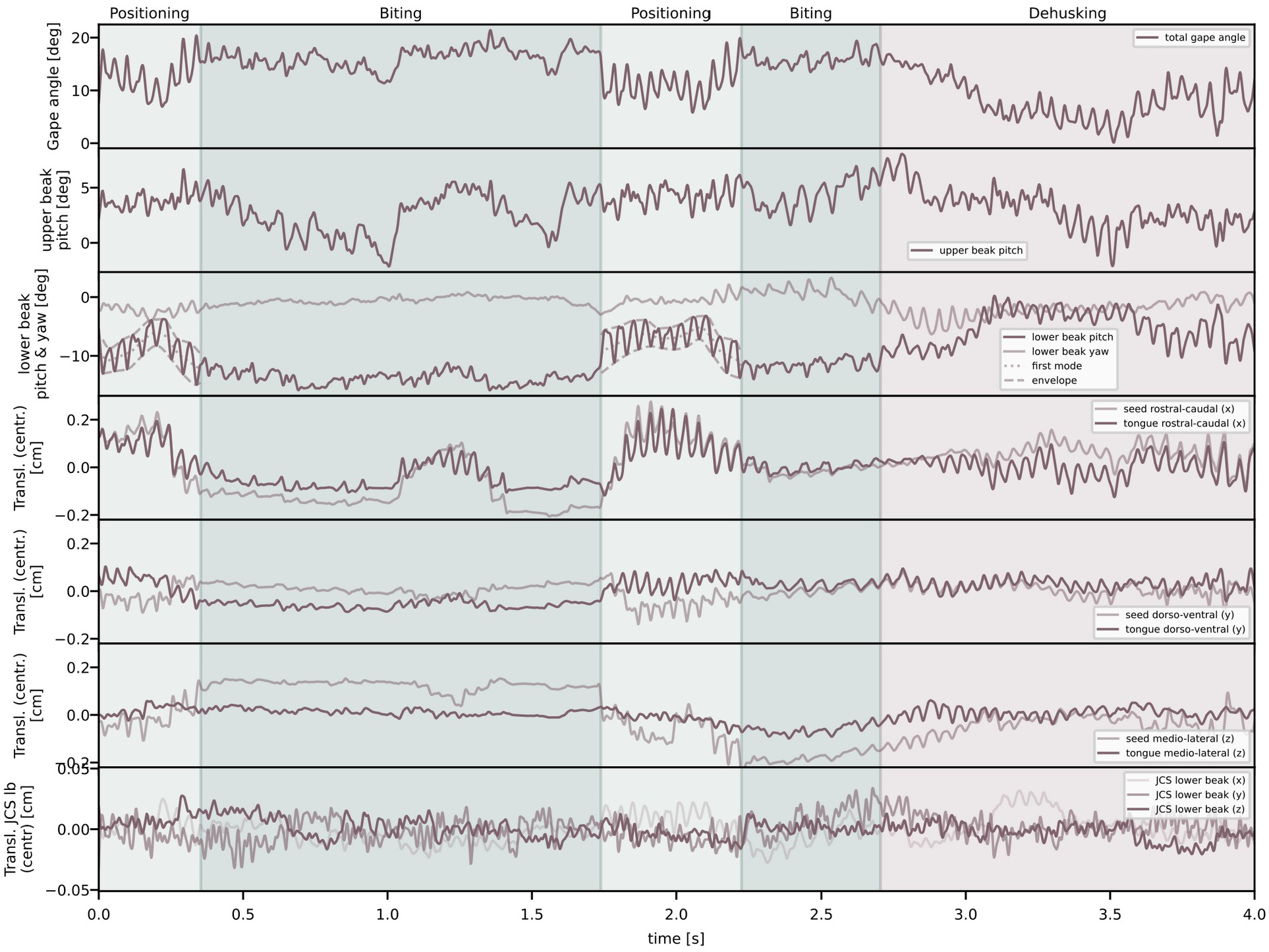
Example plots of kinematic parameters obtained from XROMM animations for a canary. From top to bottom, the panels show the following kinematic parameters as function of time: total gape pitch angle of the beak, the pitch angle of the upper beak, the pitch and yaw angle of the lower beak, the antero-posterior translation of seed and tongue, the dorso-ventral translation of seed and tongue, the medio-lateral translation of seed and tongue, and the translation of the lower beak JCS in all three dimensions. Colored shading in the background indicates the different phase types as labeled on top of the plots. Data of parameters labeled with ’centr.’ were centered (subtraction of the mean) to allow for combined visualization of multiple parameters.

**Appendix 1—figure 2.**
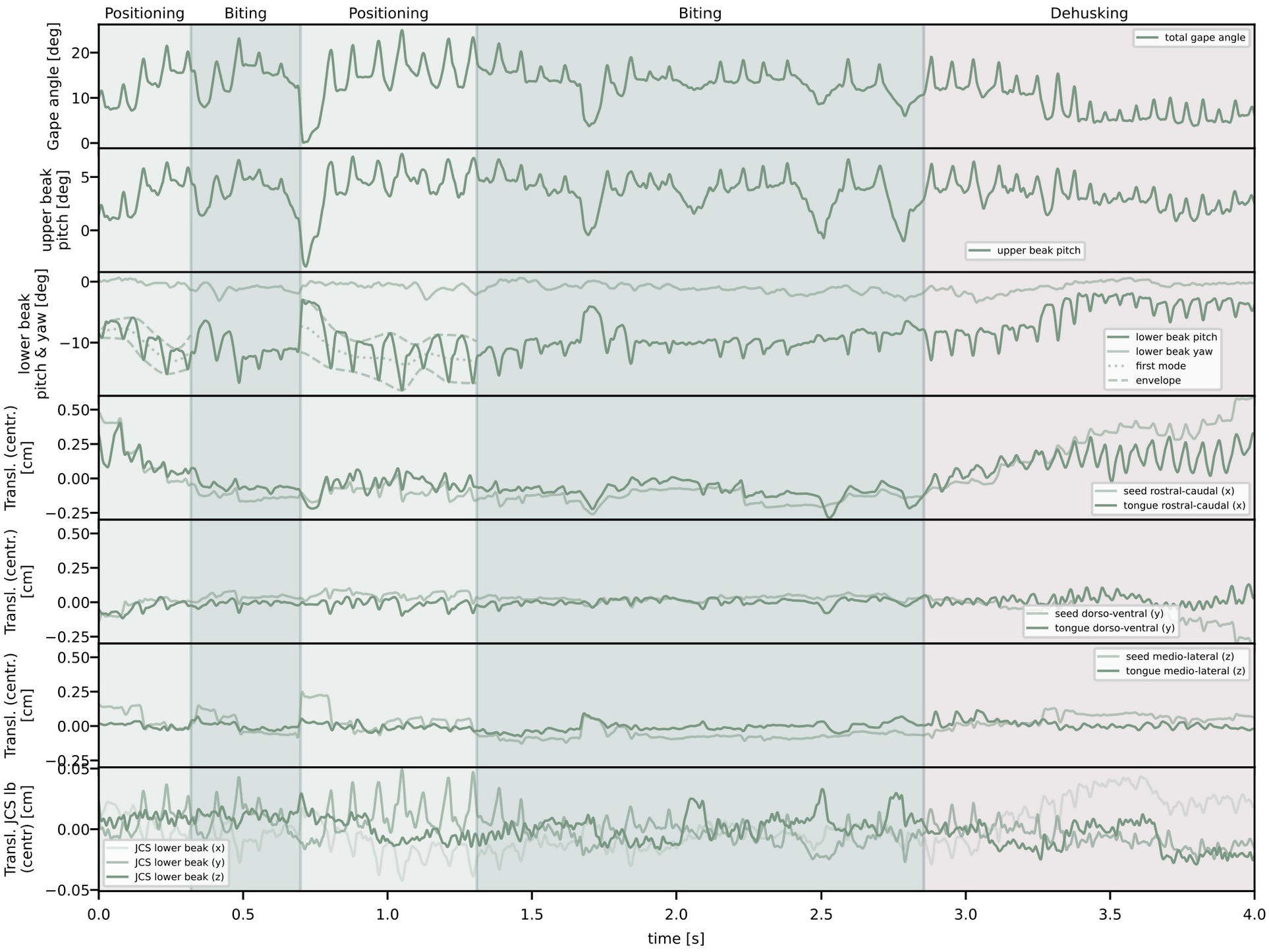
Example plots of kinematic parameters obtained from XROMM animations for a Java sparrow. From top to bottom, the panels show the following kinematic parameters as function of time: total gape pitch angle of the beak, the pitch angle of the upper beak, the pitch and yaw angle of the lower beak, the antero-posterior translation of seed and tongue, the dorso-ventral translation of seed and tongue, the medio-lateral translation of seed and tongue, and the translation of the lower beak JCS in all three dimensions. Colored shading in the background indicates the different phase types as labeled on top of the plots. Data of parameters labeled with ’centr.’ were centered (subtraction of the mean) to allow for combined visualization of multiple parameters.

**Appendix 1—figure 3.**
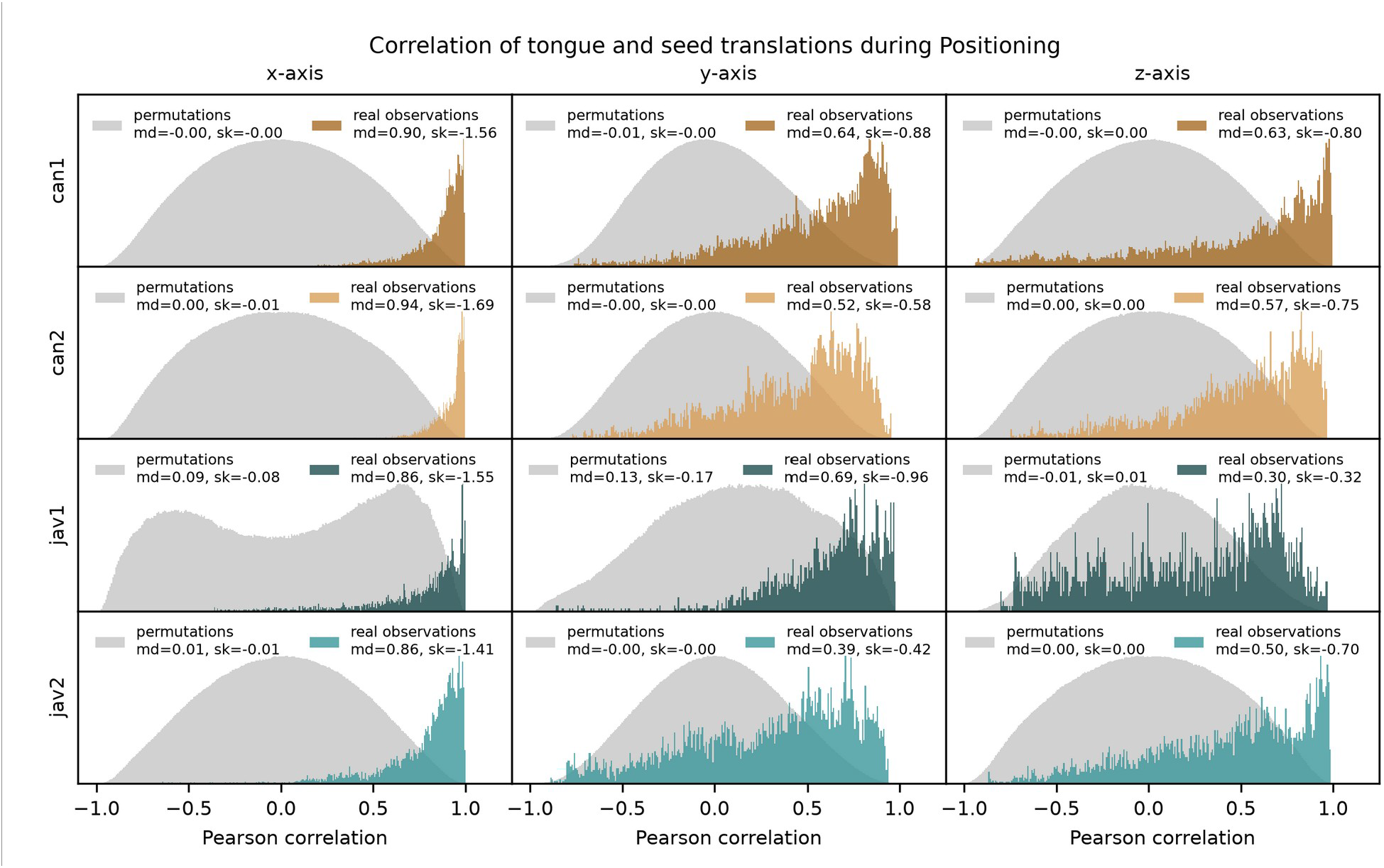
Results of the permutation tests conducted to analyze the correlation between translations of the seed and tongue markers during seed positioning. Columns relate to the three dimensions of the anatomical coordinate system (x-axis: antero-posterior; y-axis: dorso-ventral; z-axis: medio-lateral). Gray histograms show the Pearson correlation coefficients of the permuted data sequences, and colored histograms show the Pearson correlation coefficients of the real observed data. Note that in general, observed data are skewed to positive correlations, while permuted data lack a clear correlation signal. The parameter ‘md’ indicates the median of the distributions, and ‘sk’ indicates the skew of the Beta-distributions fitted to the data. The skew indicates how much the data deviates from a symmetrical distribution. The bimodal distribution of permuted data of individual ‘jav1’ along the x-axis (left column) is likely arising from particularly regular oscillations. With regular oscillations, slices shifted by multiples of the the full or half oscillatory period can yield accumulations of strong positive or negative correlation coefficients, respectively. See main text for details.

**Appendix 1—figure 4.**
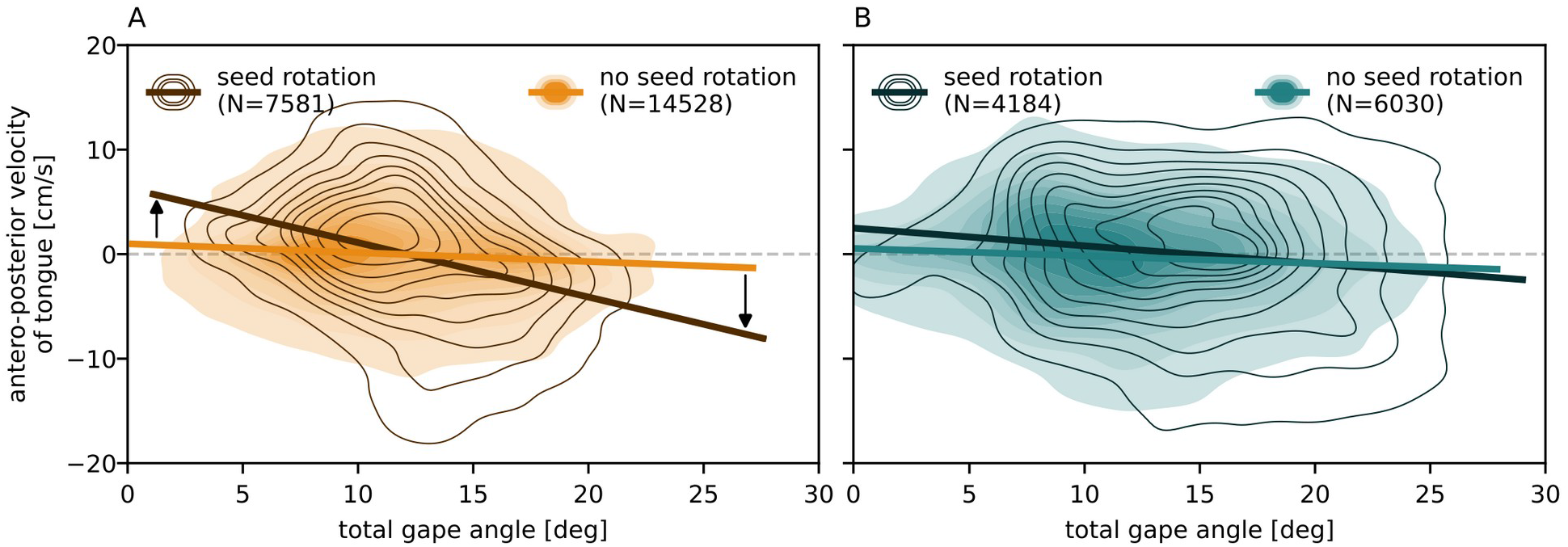
Tongue velocity during seed positioning in canaries (A) and Java sparrows (B). During seed rotation, the tongue moves faster in canaries (see black arrows in A), but much less so in Java sparrows (B). Plots show kernel density estimates of antero-posterior velocity of the tongue as function of total gape angle. More frequent combinations are indicated by darker colors. Solid lines indicate linear regressions of the data. Plots show the pooled data of two individuals per species.

**Appendix 1—figure 5.**
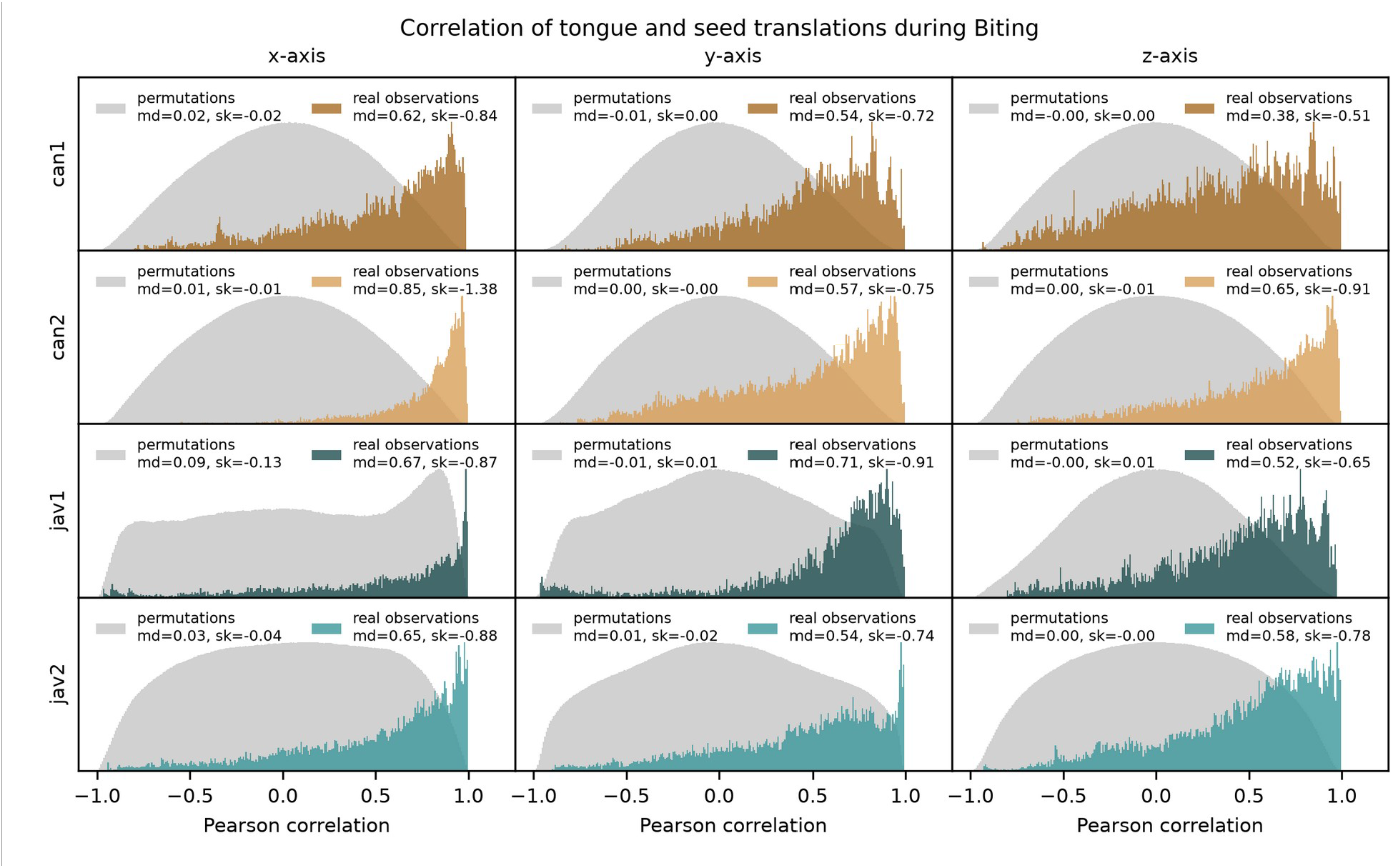
Results of the permutation tests conducted to analyze the correlation between translations of the seed and tongue markers during biting. Columns relate to the three dimensions of the anatomical coordinate system (x-axis: antero-posterior; y-axis: dorso-ventral; z-axis: medio-lateral). Labels and symbology as in Appendix 1—figure 3. Note that in general, observed data are skewed to positive correlations, while most permuted data lack a clear correlation signal. The shifted peak in the distribution of permuted data of individual ‘jav1’ along the x-axis (left column) is likely arising from a non-stationarity in the data. For example, a universal trend of anterior (or posterior) seed translation across biting trials can inflate positive correlation signals of the randomized data. See main text for details.

**Appendix 1—figure 6.**
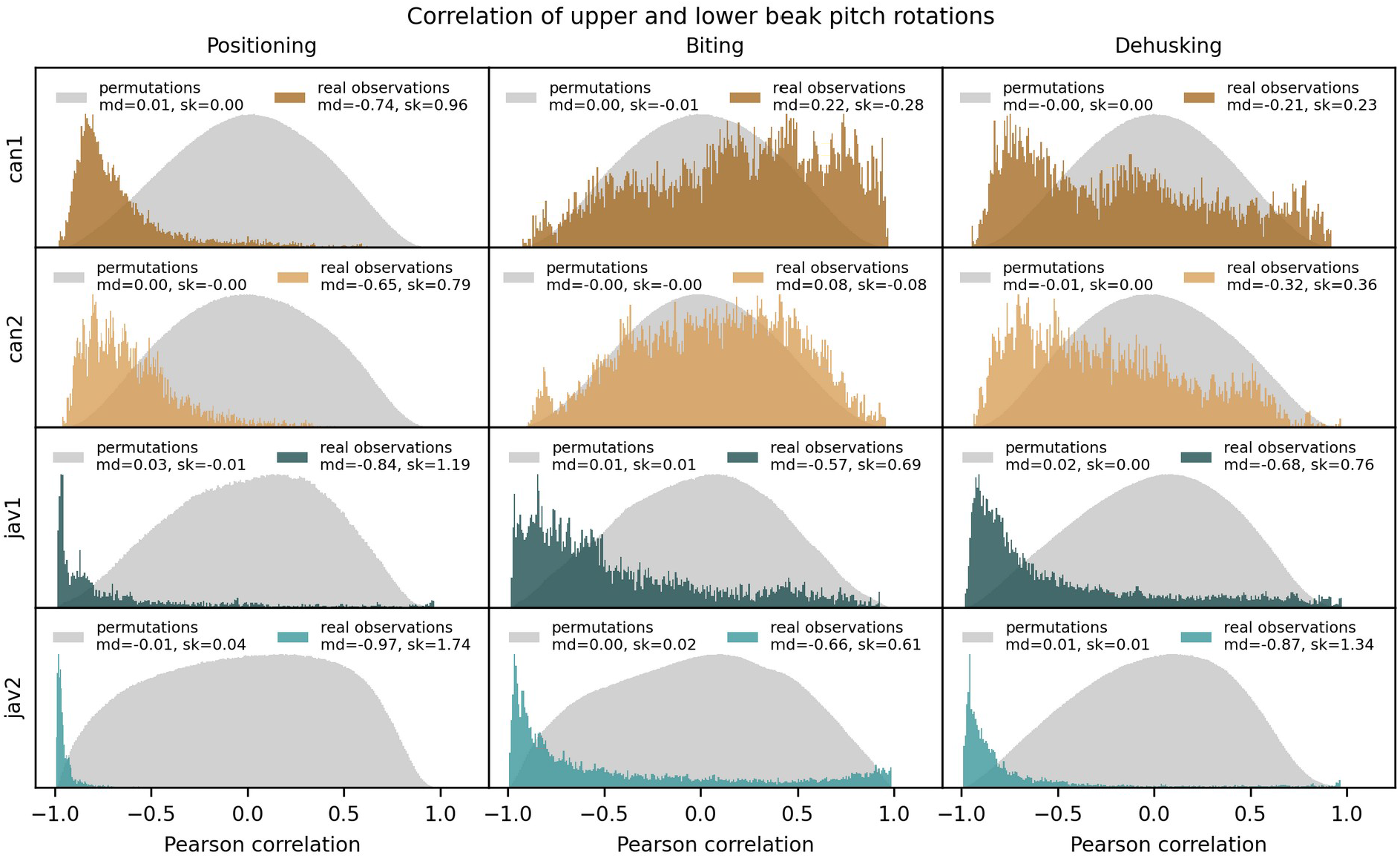
Results of the permutation tests conducted to analyze the correlation between upper and lower beak pitch rotations. Labels and symbology as in Appendix 1—figure 3. Note that permuted data show no clear correlation signal, but most observed data are skewed to negative correlations, especially during positioning, and more so in Java sparrows than in canaries. See main text for details.

**Appendix 1—figure 7.**
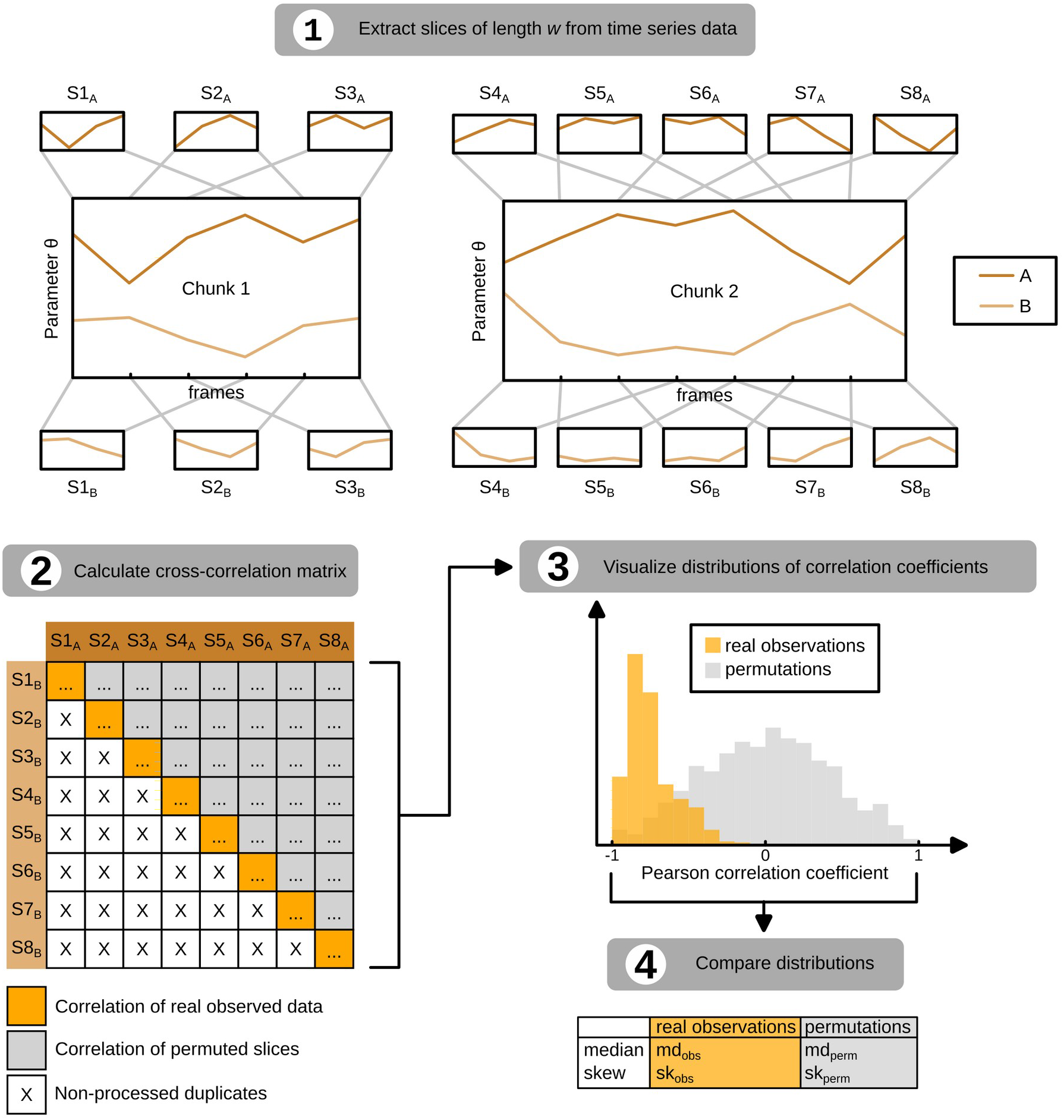
Visualization of the permutation test applied to analyze the correlation between kinematic data. Shown is a hypothetical example for analyzing the correlation between two time series, A and B, of a parameter θ, based on two chunks of data. Length *w* of the extracted slices was set to *w*=3 frames here for simplification but was set to *w*=65 frames in the actual analysis. The number of analyzed chunks (two in this hypothetical example) in the actual analysis varied between 14 and 68 depending on the individual and phasetype. See main text for details.

**Appendix 1—figure 8.**
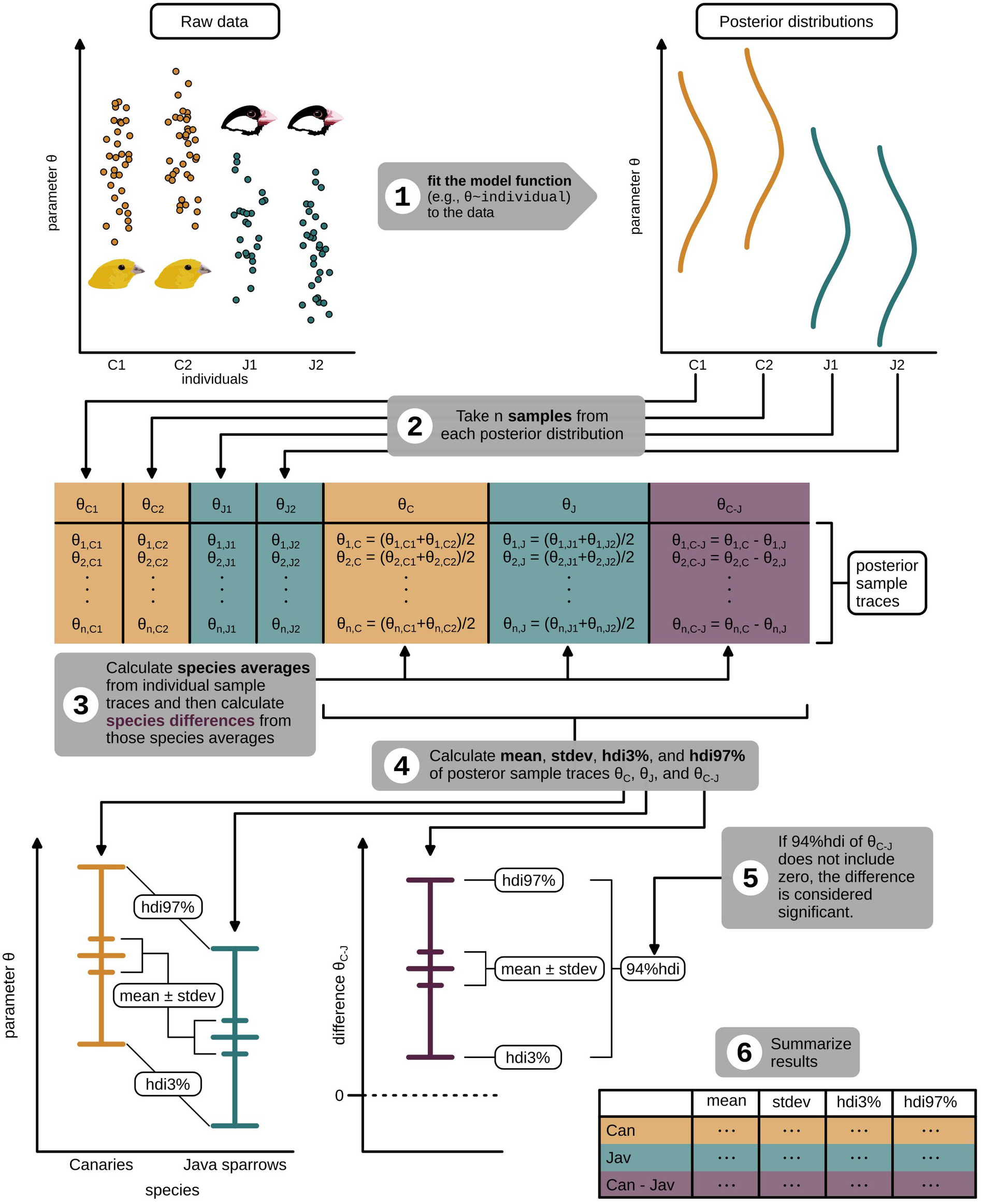
Workflow of the statistical analyses used to test for species differences (see main text for details). Plotted data are hypothetical and are for illustrative purposes only. C1, C2 and J1, J2: individual IDs of the canaries and Java sparrows, respectively; stdev: standard deviation of the mean; hdi: highest density interval. See main text for details.

**Appendix 1—table 1.** Results of the correlation analysis of the translation of seed and tongue in the three dimensions of the anatomical coordinate system. The dimensions x, y, and z refer to the antero-posterior, dorso-ventral, and medio-lateral axes of the anatomical coordinate system, respectively. The parameters ‘md’ are the medians of the distributions of Pearson correlation coefficients of the observed (obs) and permuted (perm) data. The parameters ‘sk’ indicate the skew of the Beta distributions fitted to the observed and permuted data. Note that the median and skew of permuted data are close to zero, and those of observed data are not, indicating that the correlation signal of the observed data is likely not due to chance. See main text for details.

|  |  |  | Positioning |  | Biting |  |
| --- | --- | --- | --- | --- | --- | --- |
|  |  |  | obs | perm | obs | perm |
| x-axis | Can1 | md | 0.90 | 0.00 | 0.62 | 0.02 |
|  |  | sk | -1.56 | 0.00 | -0.84 | -0.02 |
|  | Can2 | md | 0.94 | 0.00 | 0.85 | 0.01 |
|  |  | sk | -1.69 | -0.01 | -1.38 | -0.01 |
|  | Jav1 | md | 0.86 | 0.09 | 0.67 | 0.09 |
|  |  | sk | -1.55 | -0.08 | -0.87 | -0.13 |
|  | Jav2 | md | 0.86 | 0.01 | 0.65 | 0.03 |
|  |  | sk | -1.41 | -0.01 | -0.88 | -0.04 |
| y-axis | Can1 | md | 0.64 | -0.01 | 0.54 | -0.01 |
|  |  | sk | -0.88 | 0.00 | -0.72 | 0.00 |
|  | Can2 | md | 0.52 | 0.00 | 0.57 | 0.00 |
|  |  | sk | -0.58 | 0.00 | -0.75 | 0.00 |
|  | Jav1 | md | 0.69 | 0.13 | 0.71 | -0.01 |
|  |  | sk | -0.96 | -0.17 | -0.91 | 0.01 |
|  | Jav2 | md | 0.39 | 0.00 | 0.54 | 0.01 |
|  |  | sk | -0.42 | 0.00 | -0.74 | -0.02 |
| z-axis | Can1 | md | 0.63 | 0.00 | 0.38 | 0.00 |
|  |  | sk | -0.80 | 0.00 | -0.51 | 0.00 |
|  | Can2 | md | 0.57 | 0.00 | 0.65 | 0.00 |
|  |  | sk | -0.75 | 0.00 | -0.91 | -0.01 |
|  | Jav1 | md | 0.30 | -0.01 | 0.52 | 0.00 |
|  |  | sk | -0.32 | 0.01 | -0.65 | 0.01 |
|  | Jav2 | md | 0.50 | 0.00 | 0.58 | 0.00 |
|  |  | sk | -0.70 | 0.00 | -0.78 | 0.00 |

**Appendix 1—table 2.** Results of the statistical analysis for maxima in total pitch angle (cf. Figure 5A) and the corresponding pitch angles of upper and lower beak (cf. Figure 5B) for canaries (’Can’) and Java sparrows (’Jav’). Data are shown for positioning (’Pos.’), biting (’Bit.’), and dehusking (’Deh.’). Values for mean, standard deviation (stdev), and the lower and upper bounds of the highest density interval (hdi3% and hdi97%) are derived from the posterior

|  |  | max total gape angle [deg] |  |  |  | upper beak pitch at max total gape [deg] |  |  |  | lower beak pitch at max total gape [deg] |  |  |  |
| --- | --- | --- | --- | --- | --- | --- | --- | --- | --- | --- | --- | --- | --- |
|  |  | mean | stdev | hdi<br>3% | hdi<br>97% | mean | stdev | hdi<br>3% | hdi<br>97% | mean | stdev | hdi<br>3% | hdi<br>97% |
| Pos. | Can. | 15.0 | 0.1 | 14.8 | 15.3 | 4.7 | 0.1 | 4.6 | 4.8 | -10.3 | 0.1 | -10.5 | -10.1 |
|  | Jav. | 16.7 | 0.2 | 16.3 | 17.1 | 5.1 | 0.1 | 5.0 | 5.3 | -11.5 | 0.2 | -11.9 | -11.2 |
|  | Can-Jav | -1.6 | 0.3 | **-2.1** | **-1.2** | -0.4 | 0.1 | **-0.6** | **-0.2** | 1.2 | 0.2 | **0.8** | **1.6** |
| Bit. | Can. | 17.7 | 0.1 | 17.6 | 17.9 | 4.0 | 0.1 | 3.9 | 4.1 | -13.7 | 0.1 | -13.8 | -13.6 |
|  | Jav. | 16.6 | 0.1 | 16.3 | 16.8 | 4.3 | 0.1 | 4.1 | 4.4 | -12.3 | 0.1 | -12.5 | -12.1 |
|  | Can-Jav | 1.1 | 0.2 | **0.8** | **1.4** | -0.3 | 0.1 | -0.5 | 0.0 | -1.4 | 0.1 | **-1.6** | **-1.2** |
| Deh. | Can. | 13.6 | 0.1 | 13.3 | 13.8 | 4.1 | 0.1 | 4.0 | 4.2 | -9.4 | 0.1 | -9.6 | -9.2 |
|  | Jav. | 9.9 | 0.1 | 9.6 | 10.2 | 3.1 | 0.1 | 3.0 | 3.2 | -6.8 | 0.1 | -7.0 | -6.6 |
|  | Can-Jav | 3.6 | 0.2 | **3.3** | **4.0** | 1.0 | 0.1 | **0.9** | **1.2** | -2.6 | 0.1 | **-2.9** | **-2.3** |

**Appendix 1—table 3.** Results of the statistical analysis of medio-lateral movements (yaw angles) of the lower beak (cf. Figure 5D) during positioning (’Pos.’), biting (’Bit.’), and dehusking (’Deh.’). Labels ’Can.’ and ’Jav.’ refer to species names ’canary’ and ’Java sparrow’, respectively. Labels ’Can.-Jav.’ refer to differences between species. The standard deviation (’stdev’) of the distribution of used yaw angles was used to quantify the width of range of lower beak yaw rotations. Values for mean, stdev, and the lower and upper bounds of the highest density interval (hdi3% and hdi97%) are derived from the posterior distributions of the Bayesian statistical model (see Methods for details).

|  |  | mean | stdev | hdi3% | hdi97% |
| --- | --- | --- | --- | --- | --- |
| Pos. | yaw stdev. Can [deg] | 0.82 | 0.03 | 0.76 | 0.87 |
|  | yaw stdev. Jav [deg] | 0.46 | 0.03 | 0.40 | 0.53 |
|  | yaw stdev. Can-Jav [deg] | 0.35 | 0.04 | **0.27** | **0.44** |
| Bit. | yaw stdev. Can [deg] | 0.60 | 0.02 | 0.55 | 0.64 |
|  | yaw stdev. Jav [deg] | 0.57 | 0.03 | 0.52 | 0.62 |
|  | yaw stdev. Can-Jav [deg] | 0.03 | 0.04 | -0.04 | 0.10 |
| Deh. | yaw stdev. Can [deg] | 1.46 | 0.10 | 1.27 | 1.64 |
|  | yaw stdev. Jav [deg] | 0.79 | 0.07 | 0.65 | 0.93 |
|  | yaw stdev. Can-Jav [deg] | 0.67 | 0.12 | **0.43** | **0.89** |

**Appendix 1—table 4.** Results of the correlation analysis of the pitch rotation of the upper and lower beaks during positioning, biting, and dehusking. The parameters ‘md’ are the medians of the distributions of Pearson correlation coefficients of the observed (obs) and permuted (perm) data. The parameters ‘sk’ indicate the skew of the Beta distributions fitted to the observed and permuted data. The skew indicates how much the data deviates from a symmetrical distribution. Note that the median and skew of permuted data are close to zero, but those of observed data are mostly not, indicating that the correlation signal of the observed data is likely not due to chance. See main text for details.

|  |  | Positioning |  | Biting |  | Dehusking |  |
| --- | --- | --- | --- | --- | --- | --- | --- |
|  |  | obs | perm | obs | perm | obs | perm |
| Can1 | md | -0.74 | 0.01 | 0.22 | 0.00 | -0.21 | 0.00 |
|  | sk | 0.96 | 0.00 | -0.28 | -0.01 | 0.23 | 0.00 |
| Can2 | md | -0.65 | 0.00 | 0.08 | 0.00 | -0.32 | -0.01 |
|  | sk | 0.79 | 0.00 | -0.08 | 0.00 | 0.36 | 0.00 |
| Jav1 | md | -0.84 | 0.03 | -0.57 | 0.01 | -0.68 | 0.02 |
|  | sk | 1.19 | -0.01 | 0.69 | 0.01 | 0.76 | 0.00 |
| Jav2 | md | -0.97 | -0.01 | -0.66 | 0.00 | -0.87 | 0.01 |
|  | sk | 1.74 | 0.04 | 0.61 | 0.02 | 1.34 | 0.01 |

**Appendix 1—table 5.**
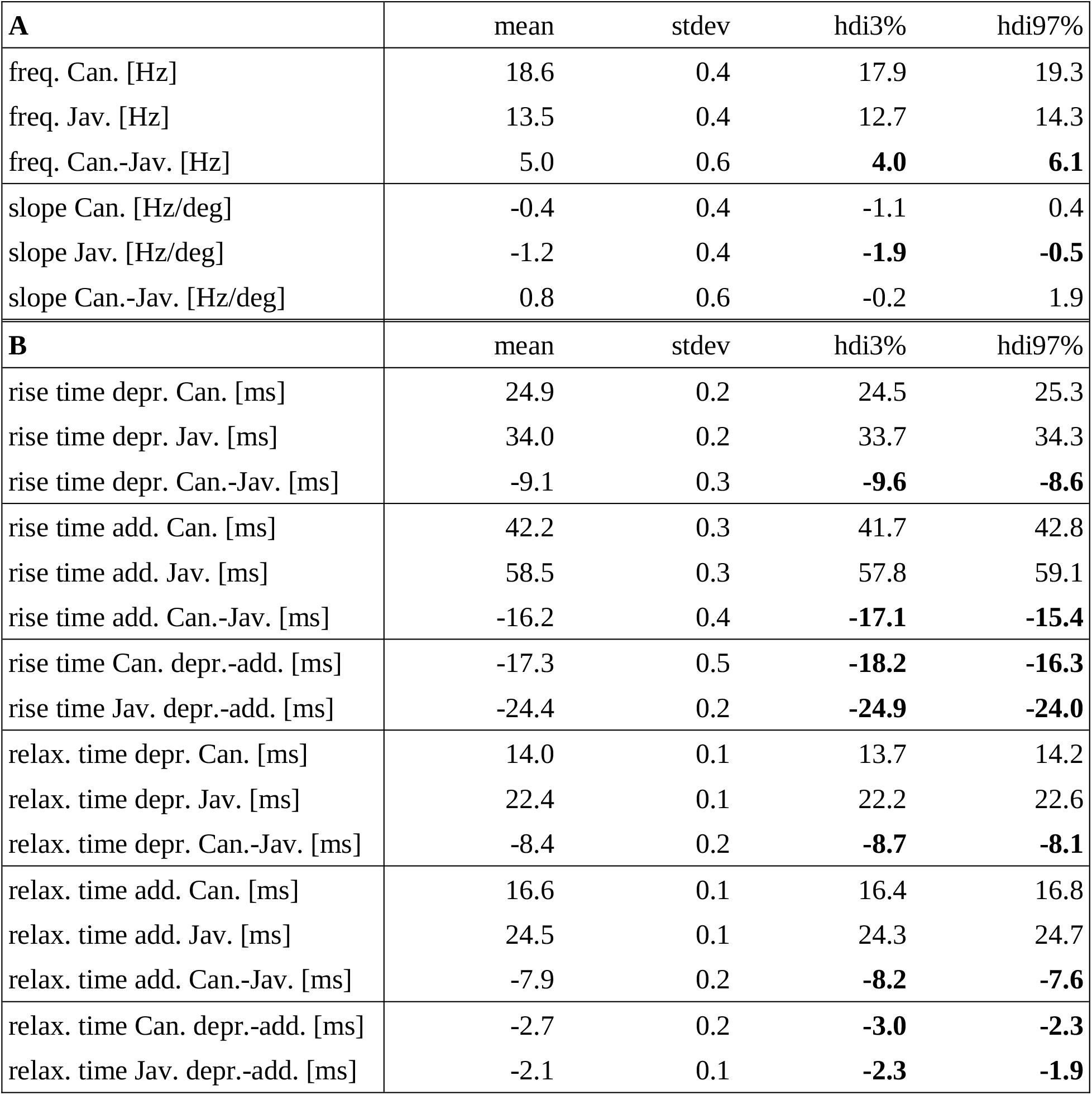
Results of the statistical analysis of A) the oscillation frequency of the lower beak (cf. Figure 8) and B) the rise and relaxation times of adductor and depressor muscles (cf. Figure 9). Labels ’Can’ and ’Jav’ refer to species names ’canary’ and ’Java sparrow’, respectively. Labels ’depr’ and ’add’ refer to the muscle types ’depressor’ and ’adductor’, respectively. Values for mean, standard deviation (stdev) and the lower and upper bounds of the highest density interval (hdi3% and hdi97%) are derived from the posterior distributions of the Bayesian statistical model (see Methods for details). Labels ’Can.- Jav.’ and ’depr.-add.’ refer to differences between species or muscle types, respectively.

